# Single-cell profiling of synchronous multi-organ metastasis reveals a systemic CD74^+^ lipid-associated macrophage niche driving polymetastatic breast cancer

**DOI:** 10.64898/2026.01.31.701004

**Authors:** Jacob Insua-Rodríguez, Pascal Naef, Hannah Savage, Isam Adam, Sung Kook Chun, Axel A. Almet, Gautham Prabhakar, Sharon Kwon, Cassandra G. Sweet, Sevilla F. Hennessey, Angela Lincy Prem Antony Samy, Hamad Alshetaiwi, Maren Pein, Sharmila Mallya, Delia F. Tifrea, Declan M. Edwards, Angela Gomez-Arboledas, Zander Esh, Carina Sandoval, Katrina T. Evans, Alex Hsu, Victoria Chen Mai, Ngoc Bao Kim Nguyen, Erika Zagni, Aaron Longworth, Paige V. Halas, Aaron Simon, Cholsoon Jang, Ahmed Mohyeldin, Robert A. Edwards, Qing Nie, Selma Masri, Kai Kessenbrock, Devon A. Lawson

**Affiliations:** Department of Biological Chemistry, University of California, Irvine; Department of Physiology and Biophysics, University of California, Irvine; Department of Mathematics, University of California, Irvine; Department of Pathology, University of Hail, Saudi Arabia; Department of Pathology and Laboratory Medicine, University of California, Irvine; Institute for Memory Impairments and Neurological Disorders, University of California, Irvine; Department of Radiation Oncology, University of California, Irvine; Department of Neurosurgery, University of California, Irvine; Department of Developmental and Cell Biology, University of California, Irvine

## Abstract

Systemic, multi-organ metastasis is the primary cause of breast cancer mortality, yet the biological mechanisms that allow disseminated tumor cells to simultaneously colonize physiologically diverse tissues remain poorly understood. Current paradigms focus on organ-specific tropism, largely overlooking the potential for systemic, conserved and synchronized programs that facilitate widespread colonization. Here, we present a high-resolution, multi-organ atlas of metastatic ecosystems and their niches using a synchronous model of brain, lung, liver, and bone metastasis combined with *in vivo* proximal niche labeling and single-cell RNA sequencing. We identify a remarkably conserved proximal niche program defined by the accumulation of CD74^+^ lipid-associated, metastasis-associated macrophages (LA-MAMs) across all metastatic sites. CD74^+^ LA-MAMs are characterized by a unique metabolic-immune signature and drive T cell suppression. We show that the cytokine Macrophage Migration Inhibitory Factor (MIF), secreted by metastatic cells, acts as the universal paracrine mediator that instructs the LA-MAM phenotype via the CD74 receptor. Interference of the MIF-CD74 axis effectively disrupts the LA-MAM niche, mitigates T cell exhaustion, and reduces metastatic burden across all organs. Analysis of a 100-patient cohort of metastasis samples from different sites confirms that the MIF-CD74 axis is a hallmark of human multi-organ colonization and independently predicts poor post-metastasis survival. Our findings define a synchronized and systemic metastatic niche that can be targeted, providing a mechanistic rationale for neutralizing the MIF-CD74 axis to treat polymetastatic breast cancer.

## Introduction

Multi-organ metastasis is the most lethal manifestation of cancer where the disease is advanced and metastatic tumors are found in multiple organs of the body. In breast cancer patients, the most common sites of distant metastasis are the brain, lung, liver, and bone^1–3^. Diagnosis of synchronous metastasis in two or more distant sites occurs in about 35% of stage IV breast cancer patients and is associated with significantly worsened outcomes as compared to metastasis restricted to one site^1,4^. Clinically, multi-organ metastasis can manifest as oligometastatic (3-5 low burden secondary tumors) or polymetastatic disease (widespread growth in multiple sites)^5^. Significant progress has been made in treatment for oligometastasis. Standard-of-care for patients with oligometastatic disease focuses on curative approaches, achieving a median survival of up to ∼100 months^6,7^. However, patients with more widespread polymetastatic breast cancer are handled with palliative care and median survival is substantially shorter (∼12 months)^8,9^. Therefore, there is a major clinical unmet need to identify new therapeutic strategies that interfere with the progression to polymetastatic breast cancer.

The prevailing “seed and soil” paradigm has traditionally focused on organotropism, identifying tissue-specific factors that make certain organs “congenial” for specific tumor types^10–13^. However, this site-specific focus fails to explain how aggressive cancer cells bypass the unique biological barriers of distinct organs simultaneously. We hypothesized that a conserved, organ-independent metastatic niche program exists, providing survival and immune evasion that supports systemic disease.

Metastatic colonization is inherently inefficient, limited by the adverse microenvironments of distant tissues^14^. To overcome this, successful disseminated cancer cells (DCCs) must rapidly instruct a supportive niche that sustains survival and favors malignant growth^15–21^. Crucially, immune evasion is a defining hallmark of the metastatic niche, acting as the ultimate gatekeeper for distant colonization^22–25^. While previous studies have utilized bulk transcriptomics or are limited to one organ, these approaches often lack the resolution to distinguish the metastatic niche from the surrounding parenchyma at multiple metastatic sites. Consequently, the universal metastatic niche drivers of polymetastatic disease have remained elusive.

To identify conserved features of metastatic niches associated with multi-organ metastasis, we developed a mouse model of synchronous metastasis to the brain, lung, liver, and bone. Using a niche-labeling system^26,27^ coupled with single-cell RNA sequencing (scRNA-seq), we systematically interrogated cellular and molecular interactions between cancer cells and their metastatic microenvironments across organs. By integrating single-cell profiling with functional perturbations and analysis of a 100-patient multi-organ cohort of metastasis samples, this study aims to define shared principles governing polymetastatic niche formation and to identify microenvironmental interactions with potential translational relevance.

## Results

### High-resolution scRNA-seq reveal cellular convergence across physiologically diverse metastatic sites

We compared the cellular and molecular composition of metastatic ecosystems and niches in brain, lung, liver, and bone using a niche-labeling system coupled with single-cell RNA-sequencing (scRNA-seq). For these studies, we developed a syngeneic mouse model of experimental metastasis using the VO-PyMT murine breast cancer cell line^28^, which generates synchronous multi-organ metastasis following injection into the left cardiac ventricle (**Figure 1a,b**). We investigated differences and similarities in the composition of the microenvironments proximal and distal to metastatic lesions using a previously established *in vivo* niche labeling system^26,27,29^. This system involves a secreted, monomeric Cherry red fluorescent protein (mCherry) containing a modified lipo-permeable Transactivator of Transcription (TATk) peptide (sLP-mCherry), and a cell-retained Green Fluorescent Protein (GFP)^26^ (**Figure 1c**). Cells engineered with this system secrete a lipid-soluble, cell membrane permeable mCherry protein (sLP-mCherry) that passively diffuses into and labels adjacent cells (**Figure 1c,d; Extended Data Fig. 1a,b**). We engineered VO-PyMT cells to express this niche labeling system (VO-sLP) and generated synchronous multi-organ metastases that we collected 12 days following intracardiac inoculation. We subsequently used fluorescence-activated cell sorting (FACS) to separate cancer cells (GFP^+^/mCherry^+^), proximal stroma cells (GFP^-^/mCherry^+^), and distal stroma cells (GFP^-^/mCherry^-^) from each organ of three different mice, encompassing 11 tissues that represented all four organs (brain, lung, liver, and bone) (**Figure 1e; Extended Data Fig. 1c; Extended Data Fig. 2a**). Cells were processed for droplet-based capture (Chromium, 10X Genomics) and scRNA-seq library preparations (**Figure 1e; Extended Data Fig. 2a,b**). scRNA-seq data analysis resulted in the identification of 73,051 cells after filtering out empty droplets and poor-quality cells (**Figure 1f; Extended Data Fig. 2b**).

**Figure 1:**
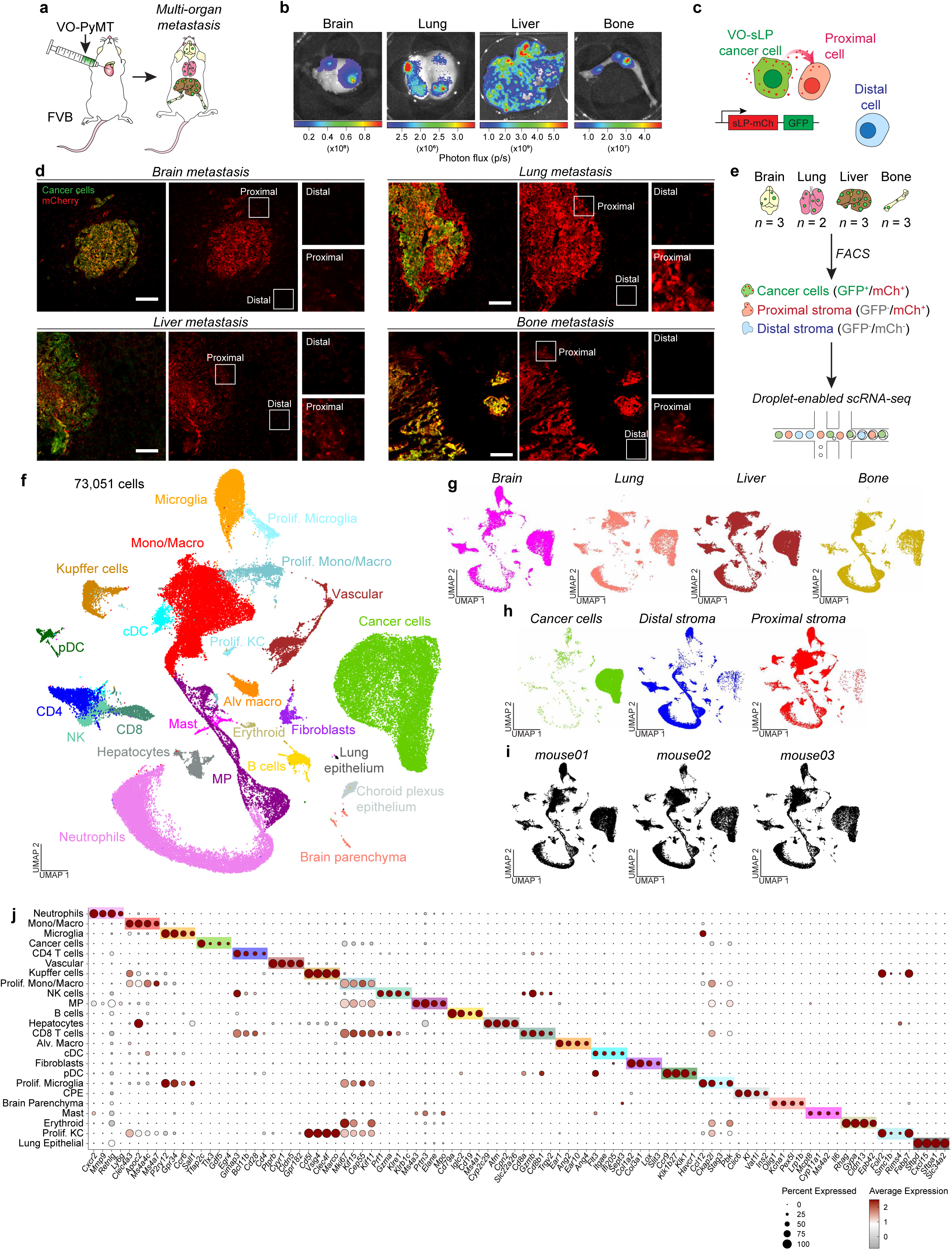
Characterization of metastatic ecosystems in brain, lung, liver and bone by niche-labeled scRNA-seq. **a.** Schematic of multi-organ metastasis model. VO-PyMT murine breast cancer cells were injected into the left ventricle of FVB female mice, and 12 days post-i.c. injection, synchronous metastases are detected in brain, lung, liver and bones. **b.** Representative *ex vivo* bioluminescence images of metastatic brain, lung, liver and lower limb bones (femora and tibiae) on day 12 post-i.c. injection of VO-PyMT cells into FVB mice. **c.** Schematic of niche-labeling system^26,27^ using GFP^+^ VO-PyMT cells (VO-sLP) secreting the lipid-soluble mCherry tag (sLP-mCh), thus labeling tumor-adjacent stromal cells. **d.** Representative micrographs showing proximal mCherry labeling of stromal cells neighboring cancer cells in brain, lung, liver and bone metastases, compared to areas distal (>40 µm) from metastatic lesions. VO-sLP cancer cells are stained with anti-GFP antibodies. Scale bar: 50 µm. **e.** Schematic of workflow used to generate a niche-labeled scRNA-seq atlas of synchronous, multi-organ breast cancer metastasis. A total of 3 brains, 2 sets of lungs, 3 livers and 3 sets of lower limb bones (tibiae and femora from both lower limbs) were mechanically and enzymatically digested into single-cell suspensions, sorted by FACS into cancer cells (GFP^+^/mCh^+^), proximal stroma (GFP^-^/mCh^+^), and distal stroma (GFP-/mCh-) populations, and subjected to droplet-enabled scRNA-seq. **f.** Uniform Manifold Approximation and Projection (UMAP) depicting 73,051 cells resulting from computational analysis of scRNA-seq data from niche-labeled multi-organ metastatic ecosystems, colored by major cell type annotations. **g.** UMAP embedding showing cells from f, split by tissue of origin. **h.** UMAP from f, split by sorted metastatic cellular compartments. **i.** UMAP from f, split by experimental replicate (i.e. mouse). **j.** Dot plot depicting average expression (circle color intensity) of 4 top marker genes for each major cell type in the scRNA-seq atlas. Circle sizes indicate the percentage of cells within each cell type expressing the indicated gene. Cell type colors from (f) are shown in the left margin.

Clustering and marker gene analysis using Seurat^30^ revealed 24 distinct cell populations (**Figure 1f,j; Extended Data Fig. 1d; Supplementary Table 1**). In addition to VO-sLP breast cancer cells (*Cldn8, Egr4, Tlx3, Epcam),* we captured numerous stromal cell types, with immune cells being the most abundant class. We captured a large frequency of myeloid cells encompassing bone marrow-derived (BMD) monocytes and monocyte-derived macrophages (Mono/Macro; *Clec4a3 , Apoc2, Ms4a4s,* Cd68), neutrophils (*Retnlg, Mmp9, Cxcr2, Ly6g)*, myeloid progenitors (MP; *Ms4a3, Prtn, Elane, Mpo*), erythroid cells (*Rhag, Gypa, Cldn13, Epb42*), mast cells (Mast; *Mcpt8, Cyp11a1, Ms4a2, Il6*), classical dendritic cells (cDC; *Flt3, Itgae, Sept3, Xcr1*) and plasmacytoid dendritic cells (pDC; *Ccr9*, *Klk1, Klk1b27*) (**Figure 1f-i; Supplementary Table 1**). We also acquired lymphoid cells including CD4 T cells (*Gimap3, Bcl11b, Cd28, Cd4*), CD8 T cells (*Cd8a, Gzmb, Cd8b1, Trgv2*), NK cells (*Prf1*, *Gzma, Klre1, Klrb1c*) and B cells (*Cd79a, Iglc2, Cd19, Ms4a1*) (**Figure 1f-i; Supplementary Table 1**). In addition to BMD immune cells, we captured tissue-resident macrophages such as microglia (*P2ry12, Gpr34, Ccr6, Tmem119*) in the brain, alveolar macrophages (Alv macro; *Ear1, Ang2, Ear10, Itgax*) in the lung, and Kupffer cells (*Clec4f, Cd5l, Vsig4, Marco*) in the liver (**Figure 1f-i; Supplementary Table 1**).

Non-immune stromal cells encompassed vascular cells (*Ptprb, Cldn5, Cyyr1, Pecam1*) and fibroblasts (*Col1a2, Col3a1, Slit3, Col1a1*). We also captured tissue-specific parenchyma such as choroid plexus epithelium (CPE) cells (*Clic6, Kl, Vat1l, Folr1*) and other parenchymal cells (*Olig1, Kcna1, Pex5l, Lrp1b*) from metastatic brains, hepatocytes from livers (Cyp2c29, *Afm, Cpn2, Alb*), and pulmonary epithelial cells from lungs (Lung epithelium; *Tinag, Sftpc, Pla2g1b*) (**Figure 1f-i; Supplementary Table 1**). At this resolution, we also detected proliferating clusters of Mono/Macro, microglia, and Kupffer cells (**Figure 1f-i; Supplementary Table 1**). These data demonstrate that we captured and characterized metastatic ecosystems and niches from diverse tissues using our experimental approach and identified convergent as well as unique cell types across anatomical sites.

### Systemic metastasis is defined by the universal enrichment of bone marrow-derived macrophages within the proximal niche

We performed subset analysis on each metastatic site to identify cell populations conserved in the proximal metastatic niches of all organs. This analysis revealed several tissue-specific cell types that were not resolved in the combined analysis, including astrocytes (*Syt10, Frmpd2, Acot5*), oligodendrocytes (*Olig1, Plp1, Ugt8a*), and neural progenitors (NP, *Sox11, Igfbpl1, Dcx*) in the brain, and classical (*Ccr2, S100a4, Ahnak*) and non-classical monocytes (*Ace, Cd300e, Adgre4*) in the bone (**Figure 2a, Extended Data Fig. 3a, Supplementary Tables 2-5**). We then analyzed the relative abundance of different cell types in the proximal and distal stroma fractions using the Milo framework, which assigns cells to partially overlapping neighborhoods within a *k*-nearest neighborhood graph^31^. Strikingly, this revealed that bone marrow derived monocytes and macrophages were consistently enriched in the proximal stroma across all four metastatic sites (**Figure 2b, Extended Data Fig. 3b**). Tissue-specific resident macrophages were also significantly enriched in the proximal stroma, including microglia in the brain, alveolar macrophages in the lung, and Kupffer cells in the liver (**Figure 2b, Extended Data Fig. 3b**). We also found enrichment of bone-specific myeloid populations in the proximal stroma, including classical (*Ccr2, S100a4, Ahnak*) and non-classical monocytes (*Ace, Cd300e, Adgre4*) (**Figure 2b, Extended Data Fig. 3b**). Vascular cells and fibroblasts (HSCs in liver, osteoblasts in bone) were the two non-immune cell types consistently more abundant in the proximal stroma (**Figure 2b, Extended Data Fig. 3b**), in line with their previously reported roles as metastatic niche components during colonization^17,18^.

**Figure 2:**
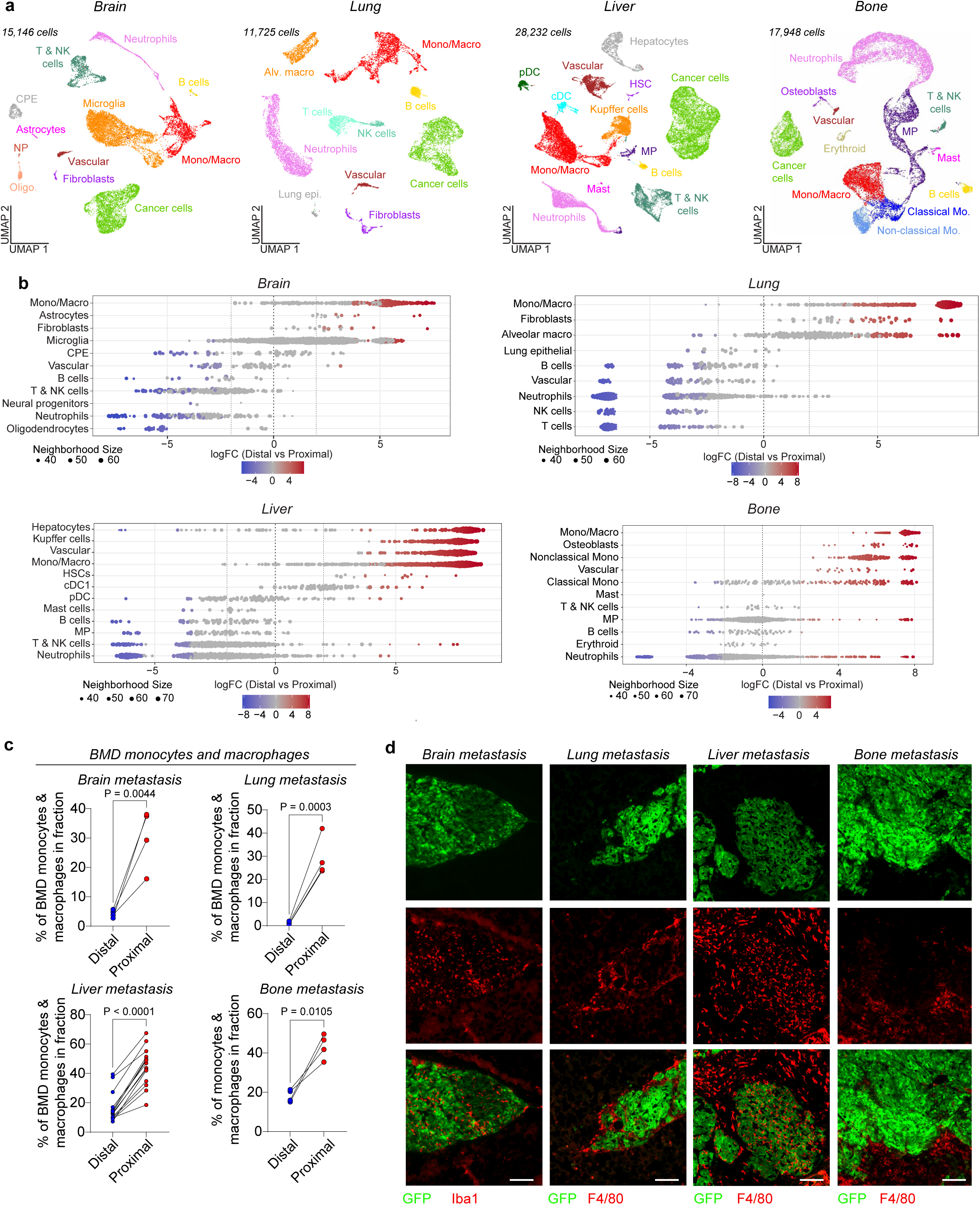
Monocyte-derived macrophages are the dominant cellular component of metastatic niches across organs. **a.** UMAPs depicting clustering of cell types in metastatic brain, lung, liver and bone tissues from FVB mice injected intracardially with VO-sLP cells. Mono/Macro, monocyte-derived macrophages; Oligo., oligodendrocytes; NP, neural progenitors; cDC1, classical dendritic cells, type 1; pDC, plasmacytoid dendritic cells; HSC, hepatic stellate cells; MP, myeloid progenitors; NK natural killer cells; Mast, mast cells; Classical Mo., classical monocytes; Non-classical Mo., non-classical monocytes. **b.** Differential abundance analysis (log fold change, logFC) of VO-sLP–labeled cells relative to unlabeled cells across phenotypic neighborhoods in brain, lung, liver, and bone metastases, analyzed using the MiloR framework. Each dot represents a cellular neighborhood grouped by cell type (indicated on y-axis). Positive logFC values indicate enrichment of labeled cells in the proximal tumor stroma; negative values indicate enrichment in distal regions. Colored dots represent neighborhoods with significant differential abundance (Benjamini-Hochberg adjusted P ≤ 0.1), with color intensity corresponding to logFC magnitude. **c.** Quantification of bone marrow-derived monocytes and macrophages in either distal or proximal stromal fractions from brain, lung, liver and bone tissues of FVB mice harboring VO-sLP metastases. Brain: *n* = 4; Lung: *n* = 4; Liver: *n* = 14; Bone: *n* = 4. P values were determined using paired *t* tests. **d.** Representative micrographs showing GFP (cancer cells) and macrophage markers Iba-1 or F4/80 expressions in brain, lung, liver and bone metastases of FVB mice harboring metastatic lesions by VO-PyMT cancer cells lacking the mCherry niche labeling system. Scale bar: 100 µm.

We found several cell types that were consistently more abundant in the distal stroma of all organs. B and T lymphoid cells were enriched in the distal stroma, suggesting that they were excluded from metastatic lesions (**Figure 2b, Extended Data Fig. 3b**). Of note, neutrophils were also predominantly found in the distal stroma of each organ (**Figure 2b, Extended Data Fig. 3b**), indicating that the sLP-mCherry tag is not indiscriminately taken up by all phagocytic cells, further supporting the finding that monocyte-derived macrophages are specifically enriched in the proximal niche.

We further investigated using protein-based, orthogonal methods the relative enrichment of myeloid cell subsets as well as CD4 and CD8 T cells in tumor-proximal or -distal stroma of mice injected with VO-sLP cells. Consistent with our scRNA-seq data, flow cytometric analysis showed that BMD monocytes and macrophages are enriched in the mCherry^+^ proximal stroma of each organ (**Figure 2c; Extended Data Fig. 4a, Extended Data Fig. 5a-d**). We also noted that CD8 T cells are significantly decreased in proximal stroma fractions from all metastatic sites (**Extended Data Fig. 4a; Extended Data Fig. 5e**), which is consistent with reports of T cell exclusion in primary tumors^32,33^. We also performed *in situ* immunofluorescence (IF) analysis to localize macrophages in metastatic lesions of mice transplanted with VO-PyMT cells, which is an orthogonal approach that does not rely on sLP-mCherry uptake for the spatial localization of proximal stroma cells. This confirmed a robust infiltration of macrophages in brain metastases (Iba1^+^), as well as lung, liver, and bone metastases (F4/80^+^) (**Figure 2d**). These data clearly establish that metastasis-associated macrophages (MAMs) represent a predominant and synchronized feature of the proximal metastatic niche that is conserved across tissues in multi-organ metastasis.

### The MIF-CD74 paracrine axis acts as a conserved master regulator of cancer cell-macrophage communication in multi-organ metastasis

We interrogated our scRNA-seq dataset for molecular interactions between cells across metastatic ecosystems using CellChat, a computational tool that infers intercellular communication networks from scRNA-seq data^34,35^. This revealed a remarkable number of predicted interactions in each tissue, highlighting the complexity of metastatic ecosystems and the value of the dataset as a resource to explore diverse cellular interactions linked to metastasis (**Figure 3a, Extended Data Fig. 6a, Supplementary Tables 6-9**). Given the abundance of MAMs at all four metastatic sites, we focused on identifying ligands produced by cancer cells and corresponding molecular receptors expressed by MAMs (Mono/Macro cluster) (**Figure 3b**). This analysis resulted in 24 significant conserved interactions (**Figure 3c,d**). Among these, we found that cancer cell-derived macrophage migration inhibitory factor (*Mif*) and its cognate, high affinity receptor *Cd74* on MAMs showed the highest communication score (**Figure 3d**). Although other cell types also express high levels of *Mif* receptors (B cells, astrocytes, Kupffer cells, cDC and pDC), *Mif* and *Cd74* expression levels were consistently the highest in cancer cells and MAMs in all four organs (**Extended Data Fig. 6b-d**).

**Figure 3:**
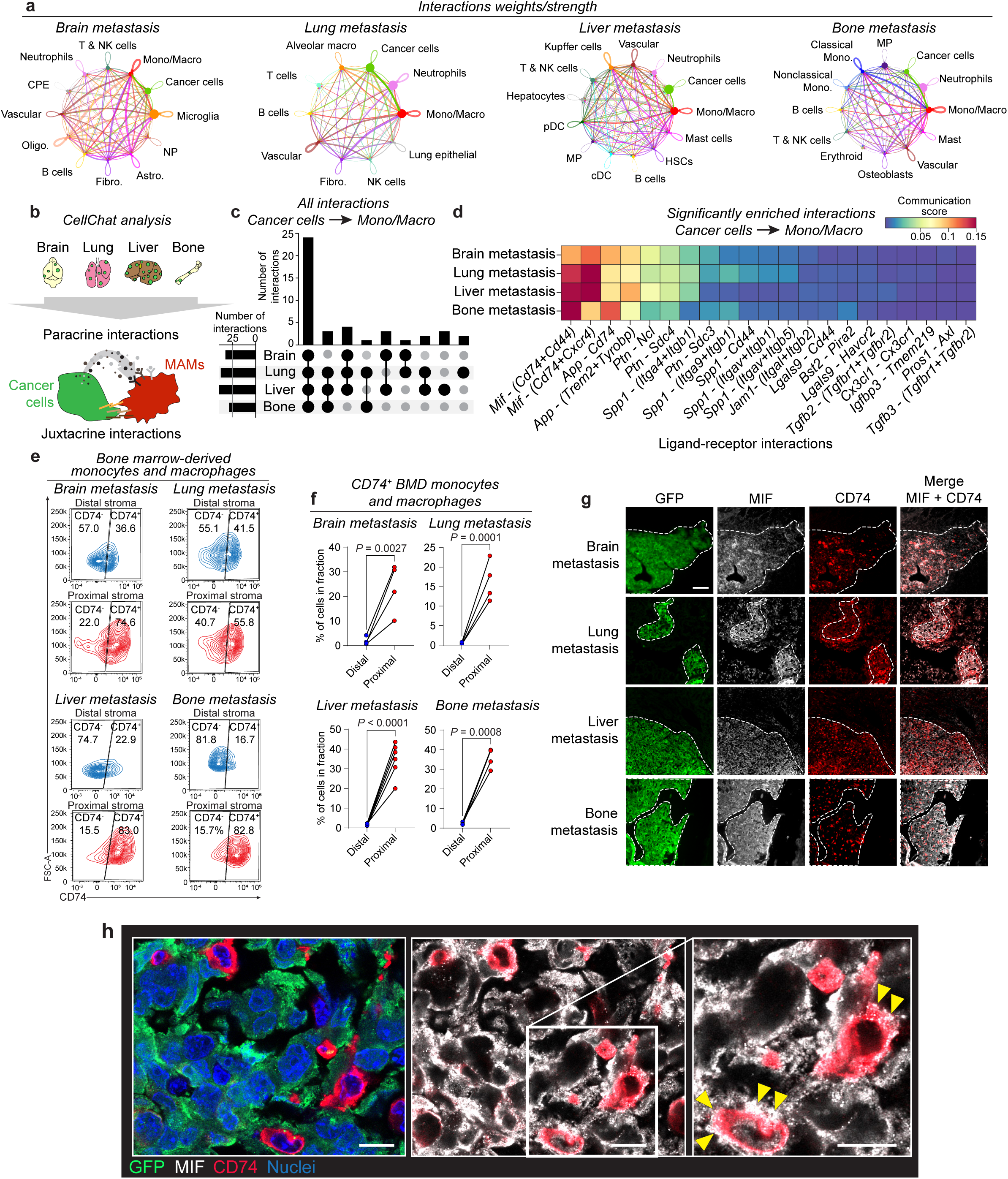
Cell interactome analysis identifies the MIF–CD74 signaling axis between cancer cells and MAMs as conserved across metastatic niches in multiple organs. **a.** Circle plots showing inferred ligand-receptor interactions among cell types in brain, lung, liver and bone metastatic ecosystems harboring VO-sLP cancer cells. Line thickness indicates interaction strength based on cell abundance and number of ligand-receptor pairs. Communication from cancer cells to monocytes/macrophages (Mono/Macro) is highlighted with opaque lines. Circle size represents relative cell-type abundance. Plots display secreted (autocrine and paracrine) and juxtacrine (cell-cell contact) interactions. **b.** Schematic of cell-to-cell communication analysis workflow using CellChat to identify paracrine interactions (ligands from cancer cells binding receptors on metastasis-associated macrophages) and juxtacrine interactions between cancer cells and MAMs in brain, lung, liver, and bone VO-sLP metastases. **c.** Upset plot showing the numbers of shared ligand-receptor interactions from cancer cells to Mono/Macro for all possible intersections between brain, lung, liver, and bone, as inferred by CellChat. **d.** Heatmap showing CellChat interaction scores of ligand-receptor interactions across all four organs. Interactions are sorted by the median interaction score. **e.** Representative contour plots showing CD74^-^ and CD74^+^ populations of monocytes and macrophages (CD45^+^/CD11b^+^/Ly6G^-^) in distal (mCherry^-^) and proximal (mCherry^+^) fractions of the indicated tissues colonized by VO-sLP breast cancer cells. **f.** Flow cytometry quantification of the relative percentage of Ly6G^-^/CD74^+^ myeloid cells (CD45^+^/CD11b^+^) relative to the total number of live cells in distal or proximal fractions of brain, lung, liver and bone tissues bearing VO-sLP metastatic lesions. Brain, n = 4; Lung, n = 4; Liver, n = 6; Bone, n = 4. P values were determined by a paired *t* test. **g.** Representative micrographs showing immunofluorescence analysis of GFP, MIF and CD74 expression in brain, lung, liver and bone tissues colonized by VO-PyMT cells lacking the niche labeling system. White dashed lines delineate metastatic lesions as determined by GFP expression in VO-PyMT cells. Scale bar: 100 μm. **h.** Confocal micrographs depicting GFP (green), MIF (white), and CD74 (red) expression in a VO-PyMT liver metastasis. Yellow arrows indicate areas where MIF signal (white puncta) accumulate in the periphery of CD74^+^ cells. Scale bars: 10 μm.

MIF is an evolutionarily conserved, small cytokine that can be secreted and is involved in several immune processes^36^. MIF binding to CD74 has been shown to trigger intracellular signaling cascades that lead to diverse cell cellular outcomes, including chemotaxis, survival, apoptosis, proliferation and transcriptional rewiring^37^. We therefore investigated the expression of Mif and Cd74 at the protein level in experimental metastases. Flow cytometry analysis confirmed a significant enrichment of Cd74^+^ BMD monocytes and macrophages (CD45^+^CD11b^+^Ly6G^int/lo^ cells) in the mCherry^+^ proximal stroma fraction as compared to the mCherry^-^ distal stroma fraction of each organ (**Figure 3e-f**). We also performed *in situ* IF analysis of Mif and Cd74 expression in VO-PyMT metastases. Consistent with our findings by flow cytometry and scRNA-seq, we found that Mif protein is highly expressed by cancer cells and Cd74 is expressed in stromal cells both within and directly adjacent to metastatic lesions (**Figure 3g**). Super-resolved confocal imaging shows that Mif co-localizes with Cd74 expressed on tumor-infiltrating, GFP^-^ stromal cells, further supporting the notion that Mif interacts with the Cd74 receptor on the cell membrane of MAMs (**Figure 3h**). These data reveal the MIF-CD74 axis as a potentially key mechanism of communication between cancer cells and MAMs that is conserved in the tumor-proximal niches across diverse metastatic sites.

### CD74^+^ MAMs adopt a conserved lipid-associated program and drive T cell suppression

Macrophages can play tumor-suppressive or tumor-promoting roles^38^. Recent single-cell transcriptomics studies have identified several recurrent subsets of tumor-associated macrophages (TAMs) that have distinct functions^39^. To investigate MAM phenotypes in higher resolution, we first performed subset analysis on myeloid cells from all four metastatic sites (**Figure 4a**). Clustering and marker gene analysis revealed nine cell populations that included cDCs, pDCs, tissue-resident macrophages, and BMD myeloid cells that separate into two lineages, monocytic and granulocytic (**Figure 4a**, **Extended Data Fig. 7a,b**, **Supplementary Table 10**). Granulocytic cells include two populations: granulocytic myeloid progenitors (GMP; *Ctsg, Elane, Mpo*) and neutrophils (*Sfta2, Ly6g, Sfta3*) (**Figure 4a**, **Extended Data Fig. 8a,b, Supplementary Table 11**). Further subclustering analysis of granulocytic cells revealed six subpopulations associated with neutrophil maturation (**Extended Data Fig. 8a-c)**. Neutrophils were predominantly enriched in the distal stroma (**Extended Data Fig. 8d-f)**, drawing clear contrast with MAMs enriched in the proximal niche.

**Figure 4:**
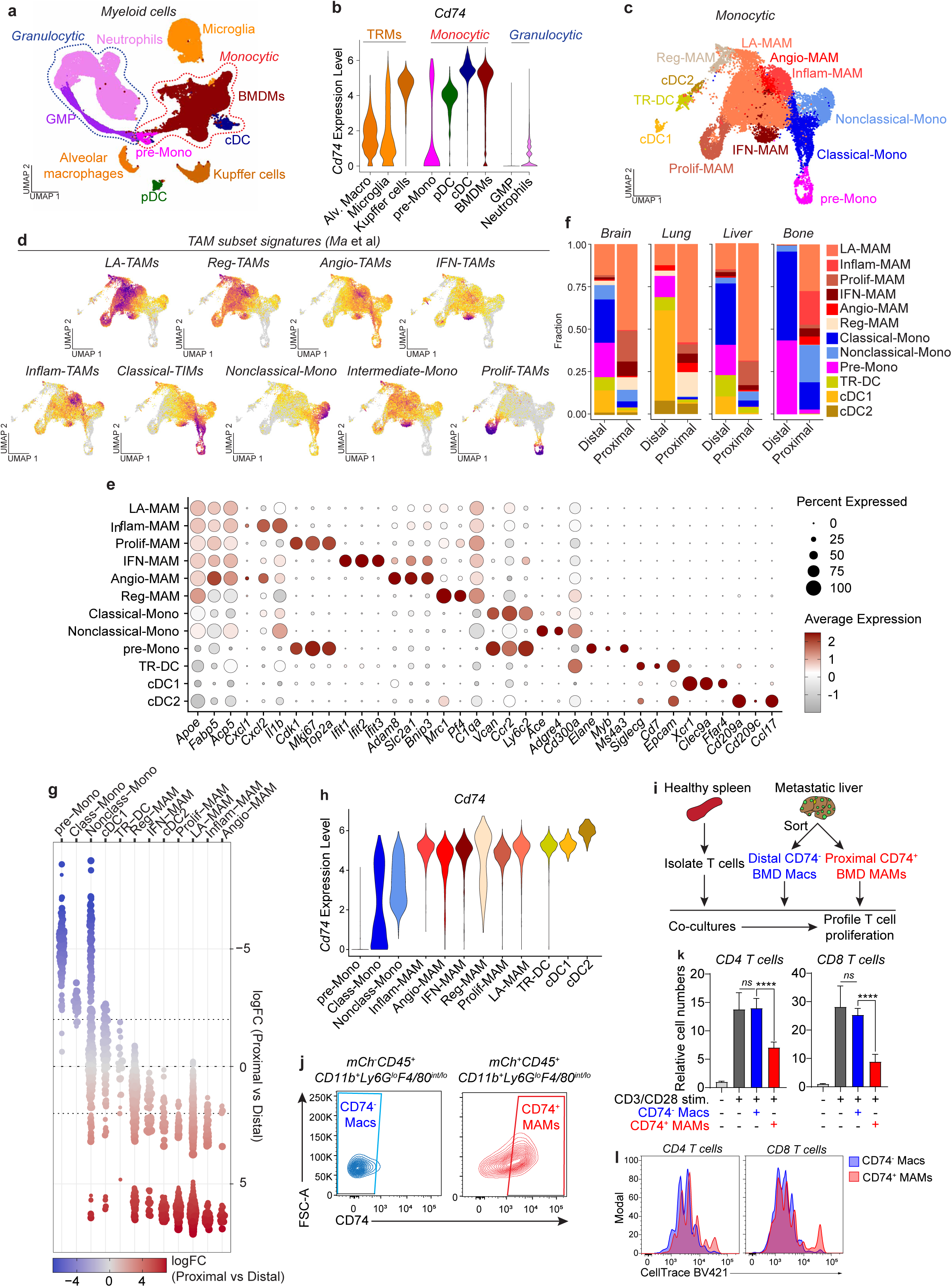
CD74⁺ MAMs exhibit a dominant lipid-associated immunosuppressive gene expression program and attenuate T cell activation. **a.** UMAP showing clustering of myeloid cells from the multi-organ scRNA-seq dataset. Cell clusters included from general object (Figure 1f) encompass: Mono/Macro, MP, Neutrophils, cDC, pDC, Microglia, Alveolar macrophages and Kupffer cells. Dashed lines enclose monocytic (pre-Mono, BMDMs and cDC) and granulocytic (GMP and Neutrophils) cellular groups. cDC, classical dendritic cells; BMDMs, bone marrow-derived monocytes and macrophages; GMP, granulocytic myeloid progenitors; pre-Mono, monocyte precursors. **b.** Relative *Cd74* expression in myeloid cells clusters, determined by Seurat. Clusters are grouped as tissue-resident macrophages (TRMs), monocytic or granulocytic clusters as indicated at the top of the plot and in (a). **c.** Subset analysis of monocytic group, encompassing Mono/Macro, cDC and pre-Mono from (a), shown as UMAP. Clusters were annotated based on previously reported TAM markers. **d.** Feature plots showing expression of the indicated gene signatures corresponding to TAM and cancer-related monocyte populations^39^ in monocytic cells captured from organs colonized by VO-sLP metastases. **e.** Dot plot heatmaps showing expression levels of 3 of the top markers for each monocytic subcluster, overlapping with corresponding, previously reported TAM subset markers^39^. Dot size represents the percentage of cells expressing the gene. Dot colors depict average expression of the gene. **f.** Relative frequencies of monocytic cell states in distal and proximal fraction of brain, lung, liver and bone tissues colonized by VO-sLP cells. **g.** Differential abundance (log fold change, logFC) of VO-sLP–labeled (proximal) monocytic cells relative to unlabeled (distal) counterparts across all phenotypic neighborhoods, grouped by the neighborhood index cell type. Positive logFC values indicate enrichment in proximal cells (red), while negative logFC indicates enrichment in distal cells (blue). Phenotypic neighborhoods with significant differential abundance (BH-adjusted P ≤ 0.1) are colored according to their corresponding logFC values. **h.** Violin plots showing *Cd74* expression levels in monocytic clusters. **i.** Schematic for T cell suppression assays using CD74^-^ bone marrow-derived macrophages (CD74^-^ BMD Macs) and CD74^+^ MAMs isolated from metastatic livers to suppress proliferation of T cells. **j.** Representative flow cytometry plots showing gating employed to sort mCherry^-^CD45^+^CD11b^+^Ly6G^-^F4/80^int/lo^CD74^-^ cells (CD74^-^ Macs) or mCherry^+^CD45^+^CD11b^+^Ly6G^-^F4/80^int/lo^CD74^+^ cells (CD74^+^ MAMs), for T cell suppression. **k.** Bar plots showing quantification of CD4 and CD8 T cell numbers upon co-culture with indicated cells. Data represents *n* = 2 independent experiments, each containing 5 technical replicates per group. *n* = 10 per group. *P* values were determined using Ordinary one-way ANOVA. *ns*, not significant; **** *P* < 0.0001. **l.** Representative flow cytometry histogram plots showing CellTrace signal in CD4^+^ and CD8^+^ T cells co-cultured with CD74^-^ macrophages or CD74^+^ MAMs.

Monocytic cells separate into three cell populations: monocyte precursors (pre-Mono; *Pif1, Itga1, Rem1*), cDCs (*Cacnb3, Ccr7, Ccl22*), and bone marrow-derived macrophages (BMDM, *Ms4a4a, Ms4a7, Arg1*) (**Figure 4a, Supplementary Table 10**). Consistent with our previous findings, we find high *Cd74* expression on myeloid cells of the monocytic lineage as well as in Kupffer cells (**Figure 4b**). Further subset analysis revealed 12 distinct cell clusters (**Figure 4c; Extended Data Fig. 7c**). These include three populations of dendritic cells: tissue-resident DC (TR-DC; *Cd7, Ppp1r14a, Ffar2*) type 1 cDC (cDC1; *Gcsam, Xcr1, Clec9a*), and type 2 cDC (cDC2; *Cd209c, Il1rl1, Ccl17*); three populations of monocytes: precursors (pre-Mono; *Pif1, Itga1, Rem1*), classical (Classical-Mono; *S1pr5, Ace, Pglyrp1*) and non-classical (Nonclassical-Mono; *Cd177, Mmp8, Vcan*); and several subsets of MAMs (**Figure 4c; Extended Data Fig. 7c**). To investigate the MAM subsets in higher resolution, we performed gene scoring using published signatures that discriminate functionally distinct subsets of TAMs (**Figure 4c-e, Supplementary Table 12**). This identified six cell states amongst the MAMs (**Figure 4c-e; Extended Data Fig. 7c, Supplementary Table 13**)^39^. The predominant population was a subset of MAMs that display a lipid-associated gene signature (LA-MAM) (**Figure 4c-f; Extended Data Fig. 7c**). These macrophages have been shown to express high levels of lipid metabolism and oxidative phosphorylation (OXPHOS) pathways and are highly immunosuppressive^39^. Of note, we found a subset of proliferating LA-MAMs (Prolif-MAM), suggesting that LA-MAMs expand locally within metastatic lesions (**Figure 4c-f; Extended Data Fig. 7c,d, Supplementary Table 13**). We identified another immunosuppressive population of regulatory MAMs (Reg-MAM) specifically in the brain and the lungs that express high levels of classic markers of alternative (M2) activation, such as *Mrc1* (encoding for CD206) and *C1qa* (**Figure 4c-f; Extended Data Fig. 7c, Supplementary Table 13**). We also found a small population of inflammatory MAMs (Inflam-MAM) that express high levels of *Il1b* and *Cxcl12* that may be linked to immune cell recruitment and activation of adaptive immunity (**Figure 4c-f; Extended Data Fig. 7c, Supplementary Table 13**). There was another subset of MAMs that display high levels of an angiogenic TAM (Angio-TAM) gene signature (*Adam8, Slc2a1, Bnip3*) (**Figure 4c-f; Extended Data Fig. 7c, Supplementary Table 13**). Angio-TAM are known to promote tissue repair through extracellular matrix (ECM) remodeling and angiogenesis. Finally, we found a population of interferon MAMs (IFN-MAM) that express numerous interferon response genes (*Ifit1, Ifit2, Ifit3*), often indicative of response to stress and inflammation (**Figure 4c-f; Extended Data Fig. 7c, Supplementary Table 13**).

We found that MAMs were consistently enriched in the tumor-proximal stroma, whereas less mature monocytic clusters were mostly captured from distal stroma fractions (**Figure 4f,g**). This was conserved in all four organs analyzed, where LA-MAMs were the dominant subset in each metastatic site (**Figure 4f,g, Extended Data Fig. 7d**). As expected, the bone harbored greater diversity of monocytic cell states, with relatively higher numbers of macrophage precursors (**Figure 4f-g; Extended Data Fig. 7d**). Importantly, *Cd74* expression was high across all MAM subsets, whereas it was lower in the monocyte clusters (**Figure 4h**). Overall, these data show that most MAM subsets are associated with immunosuppressive and other tumor-promoting functions, and that CD74 is a unifying marker for MAMs.

Given their immunosuppressive gene expression signatures, we investigated the functional capacity of CD74^+^ MAMs to suppress T cell activation in vitro^40,41^. We isolated CD4 and CD8 T cells from the spleens of tumor-naïve FVB mice and co-cultured them with either tumor-distal (mCherry^-^) CD74^-^ BMD monocytes and macrophages or proximal (mCherry^+^) CD74^+^ MAMs isolated from livers of animals harboring VO-sLP metastases (**Figure 4i,j**). Remarkably, co-culture of T cells with CD74^+^ MAMs resulted in a significant reduction in both CD4 (∼2-fold) and CD8 (∼3-fold) T cell numbers (**Figure 4k**), which is attributed to a suppression of T cell proliferation as determined by CellTrace label retention (**Figure 4l**). These findings show that CD74^+^ MAMs display a dominant lipid-associated immunosuppressive gene expression program and impair T cell activation.

### Pharmacologic or genetic disruption of the MIF-CD74 axis impairs systemic colonization across all metastatic sites

We investigated the effect of genetic or pharmacologic targeting of Mif on multi-organ metastasis. We generated stable *Mif* knockdowns using short-hairpin RNAs (shRNA) in VO-PyMT cells and in an additional model, Py8119 murine breast cancer cells (syngeneic to C57/BL6 mice)^42^. Quantitative polymerase chain reaction (qPCR), Western blot, and enzyme-linked immunosorbent assay (ELISA) showed a knockdown efficiency of >80% using two different shRNAs and confirmed that Mif is secreted into the culture media (**Extended Data Fig. 9a-e**). To determine whether Mif affects cancer cell survival or proliferation directly via autocrine signaling, we assessed the effects of *Mif* knockdown on sphere formation *in vitro*. Importantly, *Mif* knockdown did not alter the sphere forming capacity of breast cancer cells, indicating that Mif does not influence cancer cell survival or proliferation in an autocrine manner (**Extended Data Fig. 9f,g**).

We injected cancer cells expressing either control or *Mif*-targeting shRNAs intracardially into syngeneic hosts and quantified the metastatic burden in brain, lungs, liver, and lower limb bones *ex vivo*. Remarkably, *Mif* knockdown resulted in a robust and significant decrease in metastatic colonization in all four organs (**Figure 5a,b**). We performed analogous experiments using 4-Iodo-6-phenylpyrimidine (4-IPP), a small-molecule inhibitor that interferes with MIF binding to CD74^43^. We administered six doses of 4-IPP starting on day 3 after cancer cell injection (**Figure 5c**). Consistent with MIF knockdown experiments, 4-IPP treatment resulted in a substantial impairment of metastatic colonization in brain, lung, liver, and and bone (**Figure 5d**). These data show that MIF expression by cancer cells functionally promotes multi-organ metastasis *in vivo* and present the therapeutic potential of pharmacologic targeting the MIF-CD74 axis as a treatment for multi-organ, polymetastatic disease.

**Figure 5:**
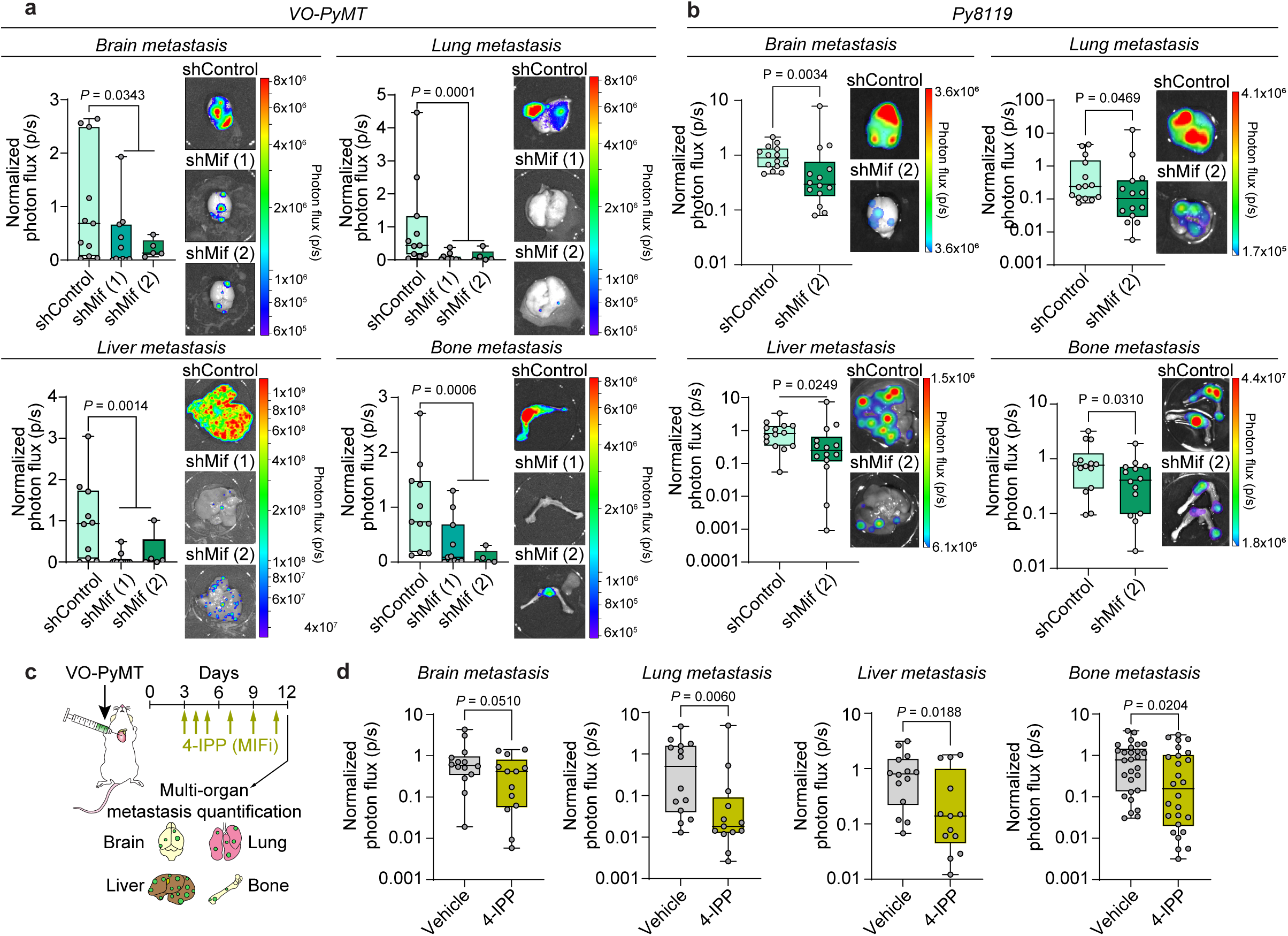
Mif targeting impairs multi-organ metastasis. **a.** Quantification of metastatic burden using *ex vivo* BLI of FVB mice injected VO-PyMT cells transduced with control vectors (shControl) and Mif-knockdown vectors (shMif 1; shMif 2). Data was acquired at day 12 post i.c. injections. shControl, *n* = 11; shMif (1), *n* = 11; shMif (2), *n* = 5. Data represents results from 2 independent experiments. In each experiment, raw photon flux values (p/s) from each organ were normalized to the average photon flux (p/s) values of the shControl group. Boxes depict 25th and 75th percentiles. Median is shown as horizontal lines. Whiskers depict data range. All data points are shown as dots. *P* values were determined using two-tailed Mann-Whitney tests by comparing shControl (*n* = 11) versus the two shMif groups combined (*n* = 16). Images show representative *ex vivo* BLI in brain, lung, liver or lower limb bones from each experimental group. **b.** Quantification of multi-organ metastatic burden *ex vivo* using BLI in albino C57/BL6 mice injected intracardially with Py8119 breast cancer cells transduced with control (shControl) and Mif-knockdown vector (shMif 2). Representative images are shown for each organ analyzed. *Ex vivo* metastatic burden was quantified 10 days after cancer cell injection. shControl, *n* = 14; shMif (2), *n* = 14. Data is representative of 3 independent experiments. Data in each organ was normalized to the average photon flux (p/s) of the shControl group in each experiment. Boxes boundaries define the interquartile ranges. Horizontal lines depict median values in each group. The whiskers show data range. The dots depict data points. *P* values were calculated by one-tailed Mann-Whitney t tests. **c.** Schematic showing pre-clinical model of MIF-CD74 targeting using 4-IPP, a small molecule inhibitor of MIF binding capacity. Multi-metastatic VO-PyMT cells were injected intracardially at day 0. Seeding and micrometastatic growth was allowed for 3 days, followed by i.p. administration of either a vehicle solution or 4-IPP. At day 12 post i.c. injection of VO-PyMT cells, metastatic colonization was analyzed *ex vivo* in brain, lung, liver and lower limb bones. **d.** Box plots showing quantification of metastatic burden of experiment as determined by *ex vivo* BLI in brain, lung, liver and bones of FVB mice injected intracardially with VO-PyMT cells and treated with the MIF inhibitor 4-IPP as described in (c). Vehicle group, *n* = 14; 4-IPP group, *n* = 13. For bone metastatic burden, right and left lower limb bones were analyzed separately. Data represents results from 2 independent experiments. In each experiment, photon flux (p/s) in each organ was normalized to the average photon flux (p/s) in the Vehicle control group. Boxes boundaries indicate 25^th^ and 75^th^ percentiles. Median is indicated by a horizontal line in boxes. Whiskers show data range. Data points are indicated as dots. *P* values were calculated using one-tailed Mann-Whitney t tests.

### MIF secretion by metastatic cells is essential for the local expansion of CD74^+^ LA-MAMs

We explored underlying mechanisms governing the prominent functional role of MIF as a multi-organ metastasis mediator. To that end, we investigated the effects of Mif attenuation on the immune microenvironment of each organ by scRNA-seq and flow cytometry. We focused on the liver for these experiments, as it is the most robust site of metastasis in this model and shows diverse immune cell populations (**Figure 2**). We also chose an early timepoint (day 7) immediately following seeding and colonization in order to reduce the confounding effects of the impaired metastatic burden that results from Mif knockdown (**Figure 5**).

We performed scRNA-seq of immune cells from livers harboring either shControl-, or shMif-expressing VO-sLP cancer cells. Immune cells were isolated by antibody-enabled magnetic activated cell separation (MACS) to enrich for CD45^+^ cells (*n* = 3 livers per group) (**Figure 6a**). Notably, this already revealed a significant decrease (1.48-fold) in the number of CD45^+^ cells in the shMif group (**Extended Data Fig. 10a**), suggesting that Mif promotes immune cell recruitment. We isolated proximal and distal stromal cells by flow cytometry using mCherry expression and subjected the cells to droplet-enabled scRNA-seq as previously described (**Figure 6a**). After removing poor quality cells and doublets, we obtained 10,955 single cell transcriptomes (*shControl* condition, *n* = 6,308; *shMif* condition, *n* = 4,647) (**Figure 6b; Extended Data Fig. 10b-d**).

**Figure 6:**
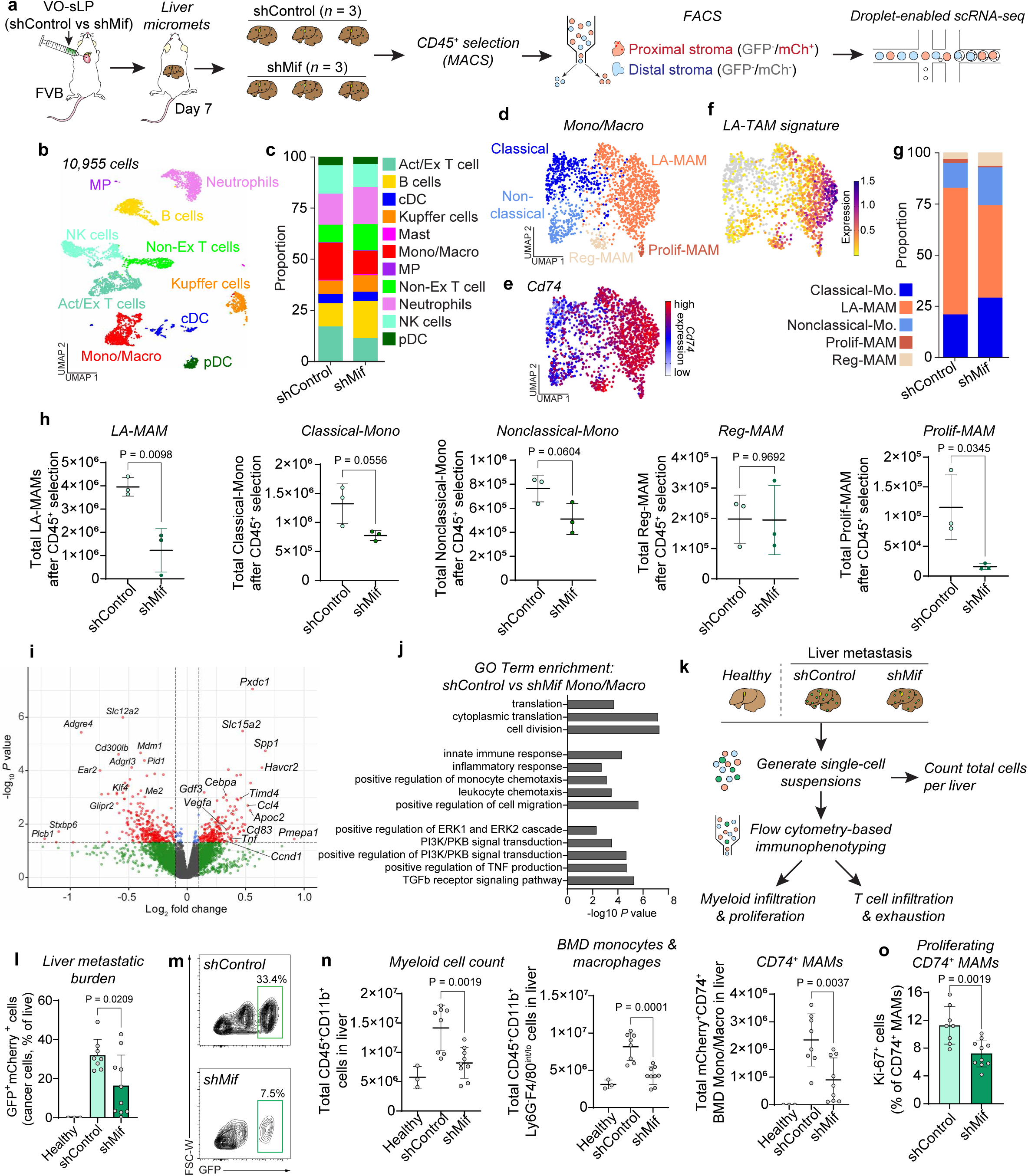
MIF promotes accumulation of immunosuppressive CD74^+^ LA-MAMs. **a.** Schematic showing strategy to characterize the effect of Mif secretion by cancer cells in the IMME. Livers with micrometastases were collected at day 7 post i.c. injection of VO-sLP multi-metastatic breast cancer cells expressing either control (shControl) or Mif-knockdown (shMif) short hairpin RNAs to attenuate Mif expression. A total of n = 3 mice per group were used. shMif (2) hairpins were used for the Mif knockdown group. Whole livers were processed for generating single-cell suspensions and CD45^+^ cells were enriched using magnetic activated cell sorting (MACS). Liver single cell suspensions were separated by FACS based on mCherry signal into distal (mCherry^-^) or proximal (mCherry^+^) fractions, and subjected to droplet-enabled scRNA-seq. **b.** UMAP depicting scRNAseq results of 10,955 cells as described in (a). Cluster names corresponding to identified cell types are indicated with the same color. **c.** Bar chart showing frequencies of annotated clusters within each group (shControl or shMif). Cell types and colors are indicated in the legend. **d.** Subset analysis of the Mono/Macro cluster from (a) shown as UMAP. LA-MAM, lipid-associated MAMs; Classical, classical monocytes; Non-classical, nonclassical monocytes; Prolif-MAM, proliferating MAMs. **e.** Feature plot showing *Cd74* expression levels in Mono/Macro cells. **f.** Module score depicting expression of the LA-TAM signature^39^ in the Mono/Macro subset depicted in (d). Relative expression levels of the signature are shown in the scale. **g.** Relative frequencies of Mono/Macro subsets in shControl or shMif groups. Subsets are indicated in the legend and correspond to the same colors as in (d). **h.** Estimated total numbers of Mono/Macro subsets calculated using the percentage of each subset out of the total liver immune cells captured by scRNA-seq analysis per mouse), and extrapolating to the total number of cells counted after whole-liver processing and CD45^+^ MACS enrichment. Horizontal lines depict mean values per group and whiskers show SD. Each data point is represented with a dot. *P* values were determined using two-tailed Student’s *t* tests. **i.** Volcano plot showing differential expression (DE) analysis of genes in Mono/Macro cells between shControl and shMif groups determined using pseudobulk analysis. X-axis depicts average log2 fold change in gene expression, while y-axis shows -log10 *P* value. A total of 17,426 genes were analyzed. Genes upregulated in the shControl group have positive log2 fold change values, whereas genes overrepresented in the shMif group have negative log2 fold change values. Dashed lines indicate cutoffs used to determine significantly regulated genes: >0.1 or <-0.1 log2 fold-change (x-axis), and *P* < 0.05 (y-axis). *P* values were determined by DESeq2. **j.** Mif-dependent GO term analysis in Mono/Macro cells. Genes upregulated in the shControl group (log2 fold-change > 0.1, *P* < 0.05) were analyzed to identify biological processes enriched in Mono/Macro cells in response to MIF using DAVID^63^. The y-axis displays selected GO terms that were significantly enriched in the shControl group. X-axis depicts *P* values of the indicated biological processes (-log10). **k.** Schematic of experimental strategy used for immunophenotyping of healthy (tumor-naive) livers and livers colonized by VO-sLP cells expressing control or *Mif*-targeting shRNAs. Whole livers were excised at experimental end point (day 10 post i.c. injections of cancer cells) and processed to generate single-cell suspensions, and total viable cell counts were determined per liver. Cells were subjected to immunophenotyping analysis by flow cytometry. **l.** Metastatic burden in livers from mice described in (k), which was determined by quantifying frequencies of GFP^+^mCherry^+^ cancer cells relative to total live cells, as determined by flow cytometry data analysis. Healthy, *n* = 3; shControl, *n* = 8; shMif, *n* = 9. Data represents two independent experiments. *P* value was calculated using two-tailed Student’s *t* test. **m.** Representative flow cytometry plots showing GFP^+^ VO-sLP cancer cells (gated in green boxes) in livers from shControl and shMif groups defined in (k). Percentages of cancer cells out of total live cells are indicated. **n.** Total counts of myeloid cell populations estimated by determining the percentage of each population out of live cells in the flow cytometry analysis, and extrapolating this percentage to the total viable cell counts from the same sample after whole liver single-cell dissociation. Each cell population is defined in the graph title, and their immunophenotype markers are defined in the y-axis titles. Healthy, *n* = 3; shControl, *n* = 8; shMif, *n* = 9. Data represents two independent experiments. *P* values were determined using two-tailed Student’s *t* test. **o.** Percentage of Ki-67^+^ cells within CD74^+^ MAMs (defined as CD45^+^CD11b^+^Ly6G^int/lo^F4/80^int/lo^mCherry^+^CD74^+^) in metastatic livers from mice injected with VO-sLP cells expressing either control or *Mif*-targeting shRNAs, as determined by flow cytometry analysis. shControl, *n* = 8; shMif, *n* = 10. Data is representative of two independent experiments. *P* values were determined using two-tailed Student’s *t* test.

Clustering and marker gene analysis revealed 11 clusters of immune cells, which included most of the same myeloid and lymphoid cell types that we observed in the liver at day 12 **(Extended Data Fig. 3a, Figure 6b)**. At this resolution, we observed a marked reduction in the frequency of the Mono/Macro cluster in the proximal stroma of the Mif knockdown condition **(Figure 6c, Extended Data Fig. 10e)**. We performed further subclustering of the Mono/Macro population to investigate changes in MAM subsets. At this timepoint (day 7), we observed five clear subsets, including Classical-Mono, Nonclassical-Mono, Reg-MAM, Prolif-MAM, and LA-MAM (**Figure 6d**). Consistent with our previous findings, LA-MAMs were characterized by high expression of *Cd74* and genes associated with the LA-TAM signature (**Figure 6e,f**). Interestingly, we found a significant reduction in the frequency of LA-MAMs in the Mif knockdown compared to control liver conditions (**Figure 6g**). We also found changes in the total numbers of MAM subsets between livers from shControl and shMif groups, which we estimated by extrapolating their relative frequencies to the total number of viable CD45^+^ immune cells counted after MACS (**Extended Data Fig. 10a**). Strikingly, Mif knockdown resulted in a significant decrease in total LA-MAMs (3.2-fold), Prolif-MAMs (7.2-fold), and a non-significant decrease in Classical-Mono and Nonclassical-Mono (**Figure 6h**). We also observed changes in gene expression in Mono/Macro upon Mif knockdown in the cancer cells. Differential gene expression and gene ontology (GO) term analysis showed downregulation of genes related to cell cycle and mitosis (e.g. *Ccnd1, Aurka, Prc1*), inflammation and leukocyte chemotaxis (e.g. *Ccl4, Cx3cr1, Ccr1*), and ERK1/2 (e.g. *Pdgfa, Vegfa, Gpr183*), PI3K (*Axl, Tnf, Osm*), and TGF-beta (e.g. *Lgals9, Tgfb3, Tgfbr1*) signaling cascades (**Figure 6i,j, Supplementary Table 14**). These data suggest that Mif promotes an expansion of immunosuppressive monocyte and macrophage subsets as well as increased immunosuppressive gene expression programs.

We further quantified differences in myeloid cell numbers in control versus Mif knockdown livers by flow cytometry. We harvested livers from mice injected with VO-sLP cells expressing either shControl or shMif hairpins, as well as tumor-naive, healthy livers (**Figure 6k**). As expected, we observed a significant reduction in metastatic burden upon Mif knockdown (**Figure 6l,m**). Mif knockdown also resulted in significantly fewer total viable cells and CD45^+^ immune cells (**Extended Data Fig. 10f,g**). Flow cytometry analysis revealed a significant reduction of total myeloid cells (CD45^+^CD11b^+,^ 1.73-fold) (**Figure 6n**), but no significant changes in Kupffer cells (Ly6G^-^F4/80^hi^) or neutrophils (Ly6G^hi^) (**Extended Data Fig. 10g**). However, Mif knockdown resulted in a significant reduction in BMD monocytes and macrophages (F4/80^int/lo^Ly6G^int/lo^, 1.9-fold) and DCs (F4/80^lo^CD11c^hi^MHC-II^hi^, 1.6-fold) (**Figure 6n; Extended Data Fig. 10g**). CD74^+^ MAMs (F4/80^int/lo^mCherry^+^CD74^+^) were also significantly reduced upon *Mif* knockdown (2.6-fold) (**Figure 6n**). Importantly, the frequency of Ki-67^+^ cells was significantly lower (1.6-fold) in CD74^+^ MAMs from livers in the shMif knockdown group, indicating that Mif also promotes MAM proliferation (**Figure 6o; Extended Data Fig. 10h,i**). Together, these data clearly show that Mif expression by cancer cells drives the proliferation and accumulation of CD74^+^ immunosuppressive monocytes and macrophages.

### Malignant MIF secretion orchestrates T cell exhaustion

We investigated whether the reduction of immunosuppressive MAMs observed in Mif knockdown livers is associated with increased T cell activation. Subset analysis of T cells resulted in four major clusters, including Natural killer (NK) cells, gamma delta (γδ) T cells, and two major clusters that were composed of both CD4 and CD8 T cells (**Figure 7a,b**). The CD4/8 T cells clearly separated into two clusters based on their differential expression of activation and exhaustion genes (**Figure 7b**). The activated/exhausted (Act/Ex) cluster expressed higher levels of *Pdcd1*, *Ctla4*, and *Lag3* and scored higher for a gene signature linked to T cell exhaustion compared to the non-exhausted (Non-Ex) cluster (**Figure 7b,c; Supplementary Table 15**). Remarkably, livers in the Mif knockdown condition harbored 2.2-fold fewern total Act/Ex T cells compared to livers from the control group (**Figure 7d,e**), suggesting that Mif promotes T cell exhaustion.

**Figure 7:**
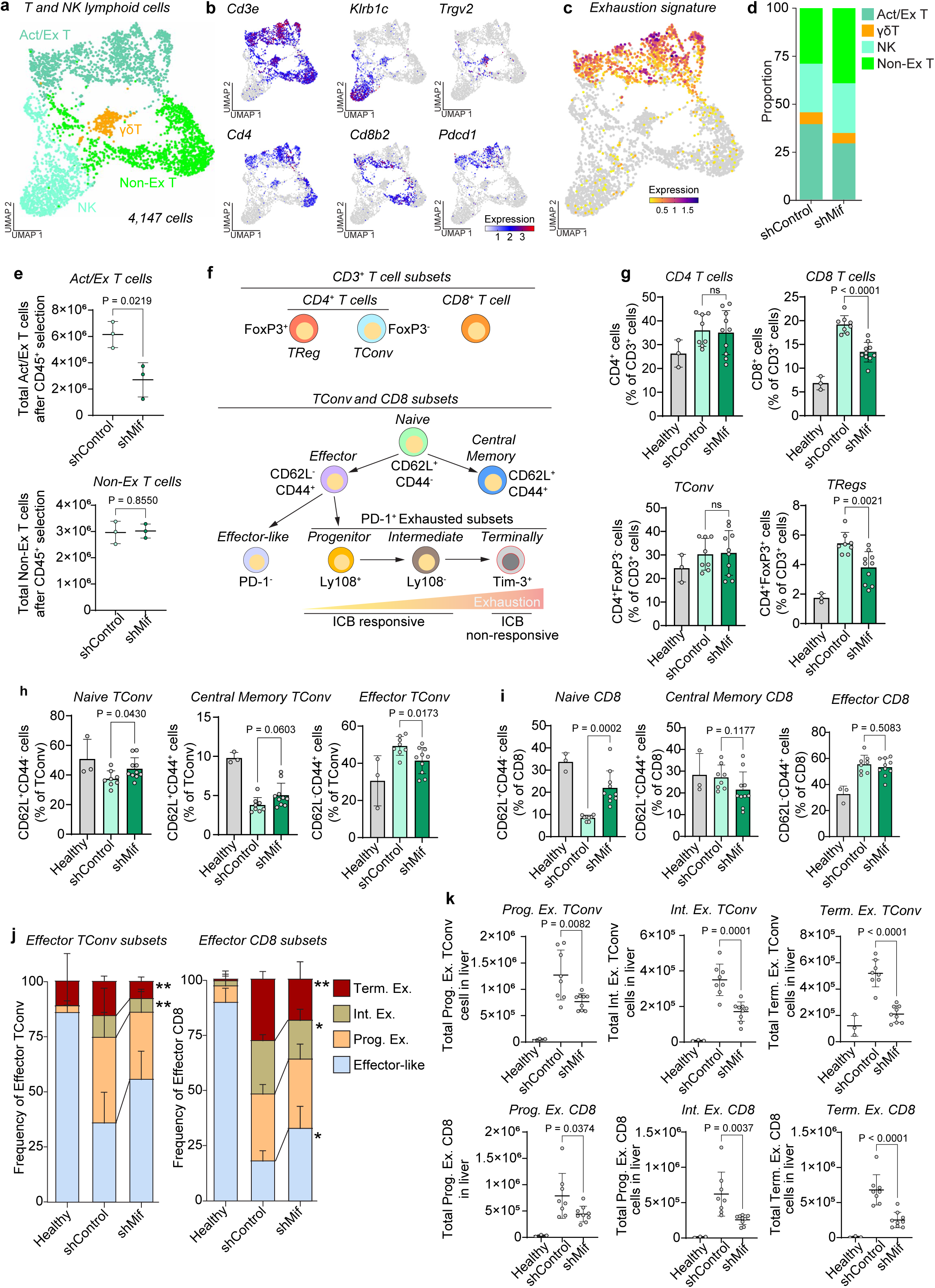
Malignant MIF secretion drives T cell exhaustion. **a.** UMAP showing subset analysis of lymphocyte cell populations (Naive T cells, Act/Ex T cells, NK cells) from scRNA-seq data (Fig. 6b). Clustering analysis using Seurat resulted in identification of an additional γδ T cell cluster (γδT). **b.** Feature plots of genes corresponding to canonical markers T cell markers: *Cd3e* (CD3) for T cells; *Klrb1c* (NK1.1) for NK cells; *Trgv2* for γδ T cells; *Cd4* for CD4 T cells; *Cd8b2* for CD8 T cells; *Pdcd1* (PD-1) for activated and exhausted T cells. **c.** Gene module scoring of T cell exhaustion gene signature expression levels in cells from (a). **d.** Relative frequencies of each lymphoid cell cluster in either shControl or shMif groups. Frequencies were calculated per group out of the total numbers in the scRNA-seq object shown in (a). **e.** Quantification of estimated total numbers of either Act/Ex T cells or Non-Ex T cells by calculating frequencies in the numbers of these clusters relative to the total number of immune cells captured in the scRNA-seq object (Fig. 7b), and extrapolating to the total numbers of viable cells quantified after whole liver processing for CD45^+^ cell enrichment. shControl, *n* = 3; shMif, *n* = 3. *P* values were determined using two-tailed Student’s *t* tests. **f.** Schematic depicting T cell populations analyzed by flow cytometry-based immunophenotyping of mouse livers in response to *Mif*-expressing or *Mif-*deficient VO-sLP metastases. T cells (CD3^+^) were subdivided as CD4 T cells (CD4^+^), and CD8 T cells (CD8^+^). CD4 T cells were further subclassified into conventional T cells (TConv, FoxP3^-^) and regulatory T cells (TReg, FoxP3^+^). TConv and CD8 cells were further classified as Naive (CD62L^+^CD44^-^), central memory (CD62L^+^CD44^+^) or effector (CD62L^-^CD44^+^) cells. Effector cells were distinguished as Effector-like (PD-1^-^), Progenitor exhausted (Prog. Ex., PD-1^+^Ly108^+^), Intermediate exhausted (Int. Ex., PD-1^+^Ly108^-^) or Terminally exhausted (Term. Ex., PD-1^+^Tim-3^+^). Arrows in the schematic depict differentiation trajectories of TConv and CD8 cells. **g.** Relative frequencies of the indicated T cell populations within CD3^+^ T cells, as determined by flow cytometry analysis of liver single-cell suspensions from healthy mice (Healthy), or mice injected i.c. with VO-sLP cells expressing either control (shControl) or *Mif*-targeting (shMif) shRNAs. Healthy, *n* = 3; shControl, *n* = 8; shMif, *n* = 10. Dots represent individual data points. Bar heights depict mean values, and error bars show SD. *P* values were determined using two-tailed Student’s *t* tests. **h.** Frequencies of TConv subsets defined by CD62L and CD44 expression, as determined by flow cytometry analysis in livers from healthy (*n* = 3), shControl (*n* = 8) or shMif (*n* = 10) groups. Subsets are indicated in the graph titles, and markers used to define each subset are indicated in the y-axis titles. Dots depict individual data points. Bar heights show mean values of each group, and error bars depict SD. *P* values were determined using two-tailed Student’s *t* tests. **i.** Frequencies of CD8 subsets based on CD62L and CD44 expression, from the same samples and as analyzed in (h). **j.** Relative frequencies of either effector TConv or effector CD8 subsets based on expression of the exhaustion markers PD-1, Ly108 and Tim-3, in either healthy livers (*n* =3), or livers colonized by *Mif*-expressing (shControl, *n* = 8) or *Mif*-deficient (shMif, *n* = 10) VO-sLP murine breast cancer cells. Bar segments are colored to depict each subset. Error bars show SD. Asterisks indicate significant differences between shControl and shMif groups of the adjacent subset (bar segment). * *P* < 0.05; ** *P* < 0.01. *P* values were determined by two-tailed Student’s t tests in comparisons of each subset. **k.** Total estimated numbers of the indicated PD-1^+^ TConv and CD8 T cell subsets in either healthy, shControl or shMif groups. Data was calculated by determining the frequency of each subset out of live cells using flow cytometry analysis, and extrapolating to the total number of viable cells quantified after whole liver dissociations. Healthy group, *n* = 3; shControl group, *n* = 8; shMif group, *n* = 9. shControl and shMif groups correspond to mice injected with VO-sLP cells expressing control or *Mif*-targeting shRNAs. Horizontal lines depict mean values of each group, and error bars show SD. Each dot represents a data point from one mouse. *P* values were determined using two-tailed Student’s *t* tests.

We further quantified differences in T cell subsets in control and *Mif* knockdown livers by flow cytometry (**Figure 7f**). We used established markers for identification of CD4 and CD8 T cell subsets by flow cytometry: regulatory T cells (TRegs), T conventional (TConv), naive, central memory, and effector. Within CD4 and CD8 effectors, we further resolved effector-like, progenitor exhausted (Prog. Ex.), intermediate exhausted (Int. Ex.), and terminally exhausted (Term. Ex) subsets (**Figure 7f**)^44^. These represent progressively more exhausted T cell states that are decreasingly responsive to immunotherapy (**Figure 7f**)^44^. Remarkably, we found that Mif abrogation is associated with an overall increase in the frequency and number of T effector subsets and a decrease in immunosuppressive and exhausted subsets. Amongst CD4 T cells, we observed a significant decrease in the frequency of Tregs (CD4^+^FoxP3^+^) and a modest increase in naive (CD62L^+^CD44^-^) and central memory cells (CD62L^+^CD44^+^) (**Figure 7g,h; Extended Data Fig. 10k**). Likewise, we observed an increased frequency of naive CD8 T cells but no change in central memory cells (**Figure 7g,i; Extended Data Fig. 10k**). Most importantly, within CD4 and CD8 effector cells, we observed an increased frequency as well as number of effector-like cells (PD-1^-^), accompanied by a significant decrease in each exhausted subset, including Prog. Ex. (PD-1^+^Ly108^+^), Int. Ex. (PD-1^+^Ly108^-^), and Term. Ex (PD-1^+^Tim-3^+^) cells (**Figure 7j,k**). Together, these data clearly establish that Mif expression by cancer cells results in an increasingly immunosuppressed immune microenvironment characterized by decreased T effector cells and increased T cell exhaustion. These findings highlight the potential of targeting MIF as a strategy to prevent or reverse T cell exhaustion and sensitize multi-organ disease to immunotherapy.

### The MIF-CD74 signature is a hallmark of human polymetastasis and independently predicts poor post-metastasis survival

We explored the clinical relevance of the MIF-CD74 axis in human breast cancer. Analysis of breast tissue data (*n* = 1,211) from The Cancer Genome Atlas (TCGA)^45^ shows that *MIF* and *CD74* RNA expression are significantly elevated in primary breast tumor tissues compared to paired or unmatched healthy breast tissues (**Extended Data Fig. 11a,b**). We further explored *MIF* and *CD74* expressions in the breast cancer molecular subtypes using the METABRIC dataset (*n* = 1,980)^46^. This showed minor but significant differences in *MIF* and *CD74* expression across the subtypes, except for the Claudin-low subtype that showed distinct expression levels (**Extended Data Fig. 11c**). Interestingly, we found that *MIF* and *CD74* expression levels increase with higher tumor grade, displaying the highest levels in grade III tumors (**Extended Data Fig. 11d**). Grade III tumors are the most aggressive, and multi-organ metastatic breast cancer is tightly associated with grade III histopathology in the primary tumor diagnosis^47^. These data show that *MIF* and *CD74* expression are higher in breast tumors compared to healthy tissue and are linked to aggressive disease.

We further determined whether *MIF* and *CD74* are preferentially expressed by cancer cells and macrophages in human breast cancer metastases as we observed in mouse models. We analyzed a publicly available scRNA-seq dataset of tumor biopsies collected from primary breast tumors and six metastatic sites (liver, axilla, neck, bone, and lung) of 30 breast cancer patients^48^. This analysis confirmed that *MIF* expression is highest in cancer cells in each organ, while *CD74* is highly expressed in the macrophage and B cell clusters (**Figure 8a,b; Extended Data Fig. 11e,f**). We also investigated cell type specificity using an additional dataset where epithelial and stromal regions of breast tumors were laser-captured microdissected^49^. This further confirmed that high *MIF* expression is virtually restricted to cancer cell regions, while *CD74* is mostly associated to stromal areas (**Extended Data Fig. 11g**).

**Figure 8:**
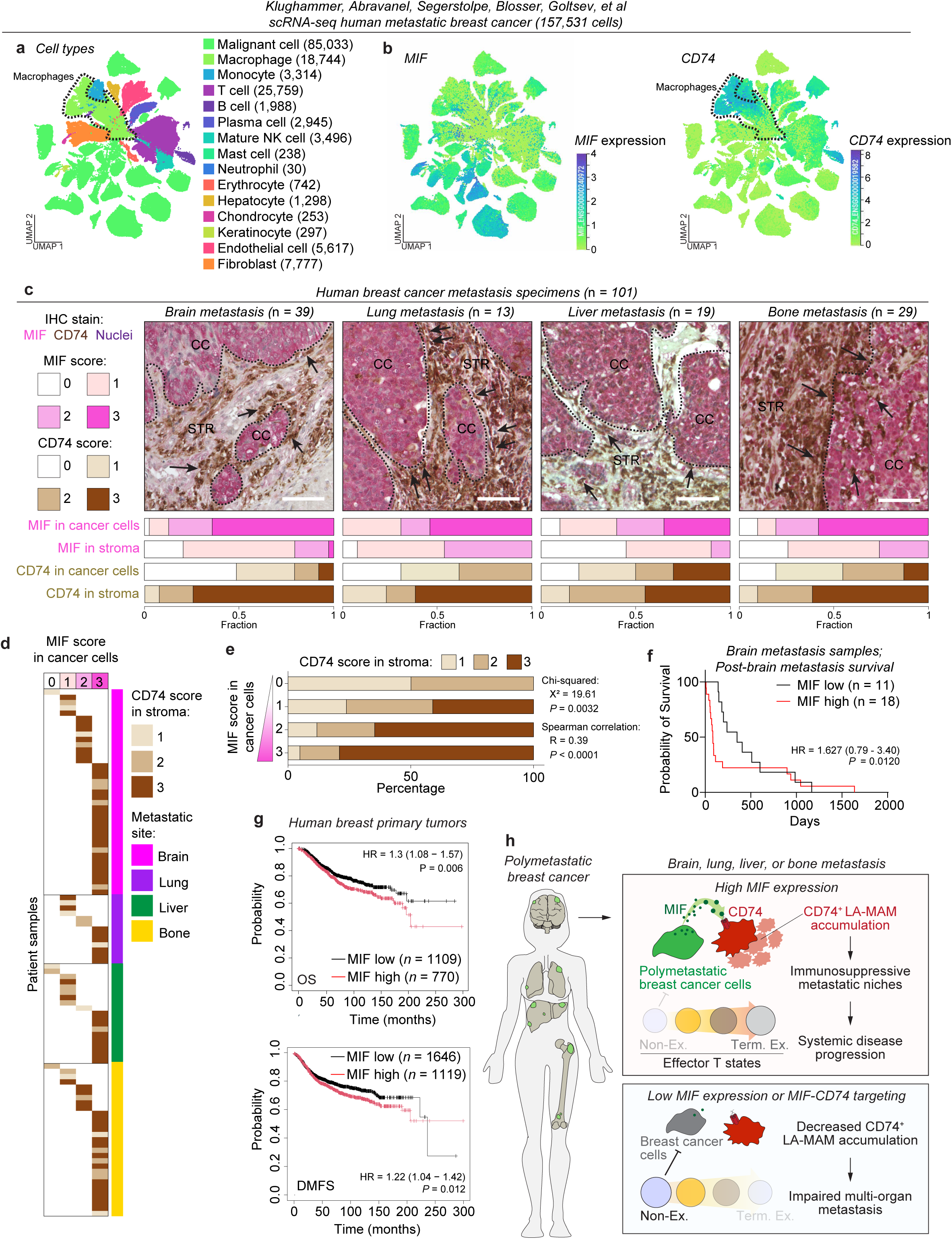
MIF-CD74 axis expression is conserved in human brain, lung, liver, and bone metastasis, and is associated with poor prognosis. **a.** UMAP showing scRNA-seq dataset of human metastatic breast cancer^48^, colored by main cell types. The number of cells per cell type are indicated in the legend. Data was collected from 6 different sites: liver, axilla, breast, bone, and lung. *n* = 30 donors. Data was accessed and plots were generated using Cellxgene^64,65^. **b.** Feature plots depicting *MIF* and *CD74* relative RNA expression. **c.** *In situ* MIF and CD74 protein expression analyzed by immunohistochemistry in a cohort of 103 human breast cancer metastases in brain, lung, liver and bone. Micrographs show representative stainings. Black dashed lines delineate areas of cancer cells (CC) in metastatic lesions. Black arrows show tumor-adjacent or tumor-infiltrating CD74^+^ stromal cells. MIF and CD74 semi-quantitative IHC scores are indicated in the color legends and frequencies of MIF and CD74 scores in either cancer cells (CC) or stroma (STR) are shown in stacked bars for each organ. Scale bar: 100 µm. **d.** Heatmap showing CD74 IHC scores in the stroma of human breast-to-brain, -lung, -liver and -bone metastasis samples, grouped by corresponding MIF IHC scores from cancer cells. **e.** Relative percentages of cases of CD74 IHC score in stroma grouped by MIF IHC score in cancer cells in all metastases analyzed. *n* = 103 samples (Brain metastases, *n* = 39; Lung metastases, *n* = 13; Liver metastases, *n* = 20; Bone metastases, *n* = 31). Chi-squared value and *P* value as calculated by Chi-squared test are shown. Spearman correlation test was calculated, and the Rho and *P* value of Spearman test are indicated. **f.** Stratification of post-brain metastasis survival in breast cancer brain metastasis samples was done based on MIF expression in cancer cell areas. MIF-high samples were determined by MIF IHC score in cancer cells of 3, while MIF-low samples were defined by MIF IHC score in cancer cells of 0-2. Post-brain metastasis survival in each patient was determined by calculating the time in days elapsed from the date of collection of the brain metastasis sample analyzed, and the date of death. MIF-high group, *n* = 11; MIF-low group, *n* = 18. Hazard ratio was calculated by a log-rank test, and *P* value was determined by a Gehan-Breslow-Wilcoxon test. **g.** Kaplan-Meier plot showing probability of overall survival (OS) or distant metastasis-free survival (DMFS) in breast cancer patients with high or low expression of MIF RNA expression in primary tumor tissues. OS: MIF high, *n* = 770; MIF low, *n* = 1109. DMFS: MIF-high, *n* = 1119; MIF low, *n* = 1646. Plots were generated using the KM plotter compiled dataset of mRNA breast cancer gene chip and visualized using the KM plotter tool^66^. **h.** Conceptual model depicting the role of MIF-CD74 paracrine interaction between cancer cells and CD74^+^ MAMs during multi-organ breast cancer metastasis. Overt, MIF-expressing breast cancer metastatic lesions are infiltrated by immunosuppressive, lipid-associated (LA) CD74^+^ MAMs, leading to immunosuppressive milieu and increased T cell exhaustion, which enables metastatic colonization at multiple distant sites, thus promoting progression to systemic, multi-metastatic disease. Targeting MIF blocks the accumulation of immunosuppressive MAMs, leading to dramatically reduced multi-organ metastasis, therefore representing a promising therapeutic avenue to prevent progression to polymetastatic disease.

To investigate the clinical relevance of MIF and CD74 protein expression in human breast cancer multi-organ metastasis, we collected 101 metastasis samples from 100 breast cancer patients (40 brain, 13 lung, 19 liver, and 29 bone) (**Supplementary Table 16**). We performed two-plex immunohistochemistry (IHC) analysis of MIF and CD74 and assigned scores for staining intensity (0-3) in regions of tissue containing cancer cells or stromal cells (**Extended Data Fig. 12**). Remarkably, we detected MIF protein expression in the cancer cells of nearly all samples analyzed (97/101, 96%) and CD74-expressing stromal cells in all samples (101/101, 100%) (**Figure 8c,d, Extended Data Fig. 11h,i**). This shows that MIF and CD74 protein expressions are highly conserved across patients and organs. MIF and CD74 expression were also preferentially observed in cancer cells and stroma, respectively (**Figure 8c, Extended Data Fig. 11h**), as we observed for RNA expression. We also noted a significant positive correlation between MIF expression in cancer cells and CD74^+^ stromal infiltration (**Figure 8d,e; Extended Data Fig. 11i**), further supporting the notion that malignant MIF secretion promotes CD74^+^ MAM accumulation.

Finally, we investigated whether MIF expression is associated with survival in breast cancer patients. Analysis of mRNA expression in primary breast tumors from a large publicly available dataset shows that high *MIF* levels are associated with shortened overall survival (OS, HR = 1.3), relapse-free survival (RFS, HR = 1.43), and distant metastasis-free survival (DMFS, HR = 1.22) (**Figure 8f; Extended Data Fig. 11j**)^50^. This supports the notion that high *MIF* expression in primary tumor cells is indicative of aggressive disease and increased metastasis, leading to shortened survival. We also investigated the relationship between MIF protein expression and survival in our 100-patient metastasis cohort. Although our cohort was only powered for analysis of brain metastasis (*n* = 40 with annotated survival data), we found that high MIF protein expression in cancer cells predicts significantly worsened post-brain metastasis survival (HR = 1.63) (**Figure 8g**), indicating that high MIF expression in metastatic tumors is associated with more aggressive disease. These data establish broad human disease relevance in breast cancer patients and support a model where MIF secretion by metastatic cells promotes the accumulation of immunosuppressive CD74^+^ LA-MAMs and T cell exhaustion, to establish an immunosuppressive niche favorable for systemic metastatic colonization (**Figure 8h**). These findings highlight the MIF-CD74 axis as a potential prognostic biomarker and new opportunity for therapeutic targeting to treat or prevent polymetastatic multi-organ metastasis.

## Discussion

Despite advances in curative treatments for breast cancer patients with localized and low burden oligometastasis, progression to polymetastatic, multi-organ metastasis remains rapidly fatal. Therefore, there is a major clinical unmet need to identify new therapeutic strategies prevent and treat the progression to polymetastatic breast cancer. The field has largely remained focused on the organ-specific paradigm of colonization, revealing distinct cancer cell-intrinsic and tumor microenvironment factors that drive tropism to specific distant sites^51^. However, whether shared biological programs enable multi-organ colonization within the same host has remained largely unexplored mainly due to the lack of appropriate model systems. Here, we identified a conserved cellular and molecular master program that governs systemic colonization across physiologically distinct tissues. By utilizing a high-resolution, synchronous multi-organ model and *in vivo* niche labeling, we have uncovered a universal metastatic niche centered on MIF-CD74 signaling and the accumulation of lipid-associated macrophages (LA-MAMs).

Current understanding of metastasis emphasizes the “seed and soil” hypothesis, highlighting how unique tissue milieus require distinct adaptations^10,51^. This is particularly relevant during early stages of colonization. However, our data suggests that while the distant tissue composition varies, the metastatic niche converges and executes a uniform program in response to growing metastatic lesions. We found that monocyte-derived macrophages are the dominant cellular constituents of the proximal niche across all organs. This cellular convergence implies that systemic metastasis may result from a coordinated biological state that can be targeted through a single molecular vulnerability. We identify the MIF-CD74 axis as the paracrine signal that fuels this conserved niche program. While MIF has been implicated in primary tumor progression and single-site metastasis^52,53^, our work establishes its role as a master coordinator of polymetastatic immunosuppression.

Crucially, our findings extend the role of MIF beyond immune cell recruitment^54^. MIF binding to CD74 receptor complexes with CD44 can trigger intracellular signaling cascades, including ERK1/2 mitogen-associated protein kinases (MAPKs), nuclear factor kappa-light-chain-enhancer of activated B cells (NF-κB), and phosphoinositide 3-kinase (PI3K), leading to diverse cellular outcomes^36^. Our data suggests that MIF directly drives the metabolic and phenotypic reprogramming of macrophages into an LA-MAM state, as well as their expansion within metastatic niches. This is significant in two ways: 1) highly proliferative metastatic cancer cells show a strong lipid avidity which in this case they may satisfy by increasing uptake of exogenous lipids coming from LA-MAMs^55^; and 2) LA-MAMs are particularly potent immunosuppressive cells that likely contribute to immune escape during metastatic progression^56–58^. By showing that MIF abrogation prevents the accumulation and proliferation of LA-MAMs, we position MIF-CD74 as a critical switch for precision reprogramming of the tumor microenvironment.

A major clinical hurdle in treating Stage IV breast cancer is the high rate of resistance to immune checkpoint blockade (ICB) in metastatic sites^59^. Our results provide a mechanistic explanation for this failure: the MIF-instructed LA-MAM niche creates a systemic T-cell exhaustion program. We observed that MIF abrogation not only reduces the number of exhausted T-cell subsets (PD-1^+^, Tim-3^+^) but also restores the frequency of effector-like cells across multiple organs. This suggests a powerful therapeutic synergy: combining MIF inhibitors (e.g. 4-IPP) with ICB may prime the systemic environment by dismantling the LA-MAM barrier, allowing T cells to effectively target multi-organ disease. This may shift the treatment goal from palliative care, currently the only option for polymetastatic patients, toward a strategy of systemic control.

We demonstrate that our findings are conserved and relevant to human disease. In our 100-patient multi-organ cohort, malignant MIF expression and CD74^+^ stromal infiltration were nearly universal features of brain, lung, liver, and bone metastases. The fact that high MIF expression predicts significantly worsened post-metastasis survival underscores the relevance of the MIF-CD74 circuit as a primary engine of systemic lethality in breast cancer.

In summary, this study provides the first integrated single-cell transcriptomic resource that captures the synchronous evolution of the metastatic niche across the brain, lung, liver, and bone. By isolating the proximal stroma through *in vivo* niche labeling, we overcome the signal dilution inherent in bulk or organ-wide single-cell analyses, revealing a conserved cellular program that governs systemic disease. We have identified the MIF-CD74 axis as a conserved, targetable vulnerability that orchestrates the systemic metastatic niche. By shifting the focus from organ-specific biology to universal systemic programs, we provide a new roadmap for the treatment of polymetastatic disease, offering a novel avenue of research towards long-term survival in patients previously limited to palliative care.

## Materials and methods

### Cell culture

VO-PyMT murine breast cancer cells, kindly provided by Zena Werb^28^, were cultured in DMEM/F12 (1:1) medium supplemented with L-Glutamine, 2.438 g/L sodium bicarbonate (Thermo Fisher, cat. no 11320033), 10% vol/vol fetal bovine serum (FBS), and 10 IU/ml penicillin and 10 µg/ml streptomycin (P/S) (Millipore Sigma, cat. no. P4333). Py8119 mouse breast cancer cells were purchased from ATCC (cat. no. CRL-3278), and cultured in F-12K medium supplemented with 2 mM L-Glutamine, 1,500 mg/L sodium bicarbonate, 10% vol/vol FBS, and P/S. HEK 293T cells were obtained from ATCC (cat. no. CRL-3216) and cultured in DMEM medium supplemented with 4.5 g/L glucose, L-glutamine, sodium pyruvate (Corning, cat. no. 10-013-CV), 10% FBS vol/vol and P/S. Cells were cultured in incubators at 37°C, 5% CO2, and 90-95% humidity (ThermoFisher Scientific). Identity and purity of all cell lines were confirmed by short tandem repeat (STR) profiling by ATCC and were routinely tested to confirm absence of mycoplasma contamination using the MycoAlert Mycoplasma Detection Kit (Lonza, cat. no. LT07-318) according to the manufacturer’s instructions.

### Generation of GFP/Luciferase-expressing cells and niche-labeling VO-PyMT cells

To identify and isolate cancer cells from animal tissues, and quantify metastatic burden, VO-PyMT cells and Py8119 cells were genetically engineered by lentiviral transduction to express a green fluorescent protein (GFP) and luciferase gene construct (pCDH-EF1a-eFFly-eGFP, Addgene Plasmid #104834). To generate niche-labeling cancer cells, VO-PyMT cells were lentivirally transduced with the pcPPT-mPGK-attR-sLPmCherry-WPRE vector (Ximbio, cat. no. 155083). Generation of niche-labeling VO-PyMT and *in vitro* and *in vivo* validation of labeling efficiency was done following the vector’s inventors protocol^27^.

### Mouse studies

All animal studies were performed in accordance with an Institutional Animal Care and Use Committee (IACUC)-approved protocol #AUP-19-051 at the University of California Irvine. For allograft studies involving VO-PyMT cells, we employed syngeneic 6- to 8-weeks-old, female, FVB mice (The Jackson Laboratory, RRID:IMSR_JAX:001800). For allograft studies with Py8119 cells, we utilized syngeneic, 6- to 8-weeks old, female, B6 albino mice (B6(Cg)-Tyrc-2J/J, The Jackson Laboratory, RRID:IMSR_JAX:000058). Mice were housed in individually ventilated cages, in an enriched environment with bedding material and under humidity and temperature control. For experimental multi-organ metastasis assays, mice were injected with 500,000 cancer cells into the systemic circulation by intracardiac (i.c.) delivery into the left ventricle using a 31G 1 ml insulin needle (BD). In FVB animals injected with VO-PyMT cells, we determined day 12 post-i.c. injection as humane endpoint before emergence of evident signs of discomfort in mice and significant detectable metastasis in brain, lung, liver and bone. In B6 albino mice injected with Py8119 cells, humane endpoint was determined at day 10 post-i.c. injection. For metastatic burden quantification, mice were sacrificed and brains, lungs, livers and tibiae and femora from both lower limbs were collected and submerged in 1.5 mg/ml D-Luciferin, Potassium Salt (Goldbio, cat. no. LUCK-100) in PBS for 10 minutes before *ex vivo* bioluminescence imaging (BLI) using an IVIS Lumina (Caliper) or an AMI-HT (Spectral Instruments Imaging) instrument.

### scRNA-seq profiling of multi-organ metastasis from mouse tissues

#### Tissue digestion for generation of single-cell suspensions

To generate a niche-labeled scRNA-seq dataset of multi-organ synchronous metastasis, single cell suspensions were generated from brain, lung, liver and bones from lower limbs tissues of FVB mice injected with VO-PyMT cells. One mouse was processed each time, and a total of 3 mice were used for this dataset. To generate single cell suspensions, tissue dissociation protocols were optimized to ensure the best possible representation of cellular diversity for each organ, cell yield and viability, considering technical feasibility as well. For each iteration of these experiments, a cohort of 3-5 mice were injected i.c. with VO-PyMT cells, and on days 10-11 post-i.c. injections, systemic metastatic burden was monitored *in vivo* by injecting 200 µl of 15 mg/ml D-Luciferin in PBS, i.p., allowing luciferase reaction for 10 minutes, and imaging mice under isoflurane anesthesia using an IVIS Lumina imager (PerkinElmer). To model high burden systemic metastasis and ensure the highest possible cancer cell and proximal stroma cell yield, one mouse with robust BLI signal from head (brain metastasis), thorax (lung metastasis), abdomen (liver metastasis) and lower limbs (bone metastasis) was selected to be processed for multi-organ scRNA-seq preparations. Liver dissociation was first performed by direct 2-step perfusion through the portal vein to ensure hepatocyte viability^60,61^. At endpoint, mice were first anesthetized using 100 mg/kg pentobarbital solution by intraperitoneal (i.p.) injection. An incision in the abdomen was made to expose the portal vein and inferior vena cava (IVC), a suture was placed around the IVC, and the portal vein was severed. Using a needle connected to a peristaltic pump by a cannula, the liver was first perfused with pre-warmed chelating buffer (25 mM HEPES, 115 mM NaCl, 5 mM KCl, 1 mM KH^2^PO^4^, 2.5 mM MgSO^4^, 0.5 mM EGTA pH 7.4) for 5 minutes, followed by perfusion with pre-warmed digestion buffer (25 mM HEPES, 115 mM NaCl, 5 mM KCl, 1 mM KH^2^PO^4^, 2 mM CaCl^2^, 0.1 mg/ml Collagenase Type IV, pH 7.4) for 15 minutes. After perfusion, gall bladder was removed and liver was collected and placed on a petri dish on ice containing liver resuspension buffer (25 mM HEPES, 115 mM NaCl, 5 mM KCl, 1 mM KH^2^PO^4^, 2.5 mM MgSO^4^, 4 mM CaCl^2^, pH 7.4) supplemented with 10 mg/ml bovine serum albumin (BSA, Fisher Scientific, cat. no. BP-9705). Livers were then peeled and gently shaken to release cells into resuspension buffer. Liver homogenates were then filtered through a 70 µm strainer and pelleted by centrifugation at 30 g for 1 minute at 4C. Supernatants contained non-hepatocyte cells (NHCs) and cell pellets were mostly hepatocytes. Supernatants were then separated, centrifuged at 300 g at 4C, and both NHCs and hepatocytes were resuspended in liver resuspension buffer supplemented with 10 mg/ml BSA. NHCs and hepatocyte suspensions were counted, mixed on a 1:1 ratio. NHCs and hepatocyte mixtures were kept on ice until cell multiplexing oligo (CMO) labeling. For generating metastatic brain single cell suspensions, brains were first chopped into 6 pieces using scalpels and simultaneous enzymatic and mechanic digestion with 2 ml 1 mg/ml Collagenase D (Millipore Sigma, cat. no. 11088858001) and 10 µg/mL Deoxyribonuclease I (Worthington Biochemical Corporation, cat. no. LS006342) in HBSS supplemented with Ca^2+^ and Mg^2+^ (HBSS^++^, Corning, cat. no. 21-023-CV) using GentleMACS C tubes (Miltenyi Biotec, cat. no. 130-093-237) and a GentleMACS Octo Dissociator with heaters (Miltenyi Biotec), for 30 minutes at 37°C using the built-in program 37C_ABDK_01. Upon enzymatic and mechanical digestion, brain homogenates were resuspended in 5 ml ice cold DMEM/F-12 medium supplemented with 10% vol/vol FBS and filtered through a 70 µm strainer (Fisherbrand cat. no. 22363548). Brain homogenates were pelleted by centrifugation at 300 g for 5 minutes at 4°C and debris was removed using 900 µl Debris Removal Solution (Miltenyi Biotec, cat. no. 130-109-398) according to the manufacturer’s instructions. 4 ml of cold HBSS^++^ was gently overlayed onto brain suspensions in Debris Removal Solution and then centrifuged at 3,000 g for 10 minutes at 4°C, yielding a triphasic solution. Two top layers (supernatant and debris) were discarded by aspiration, and bottom layer was gently resuspended in 10 ml cold HBSS^++^ by inverting three times. Brain cells were then collected by centrifugation at 1,000 g at 4C for 10 minutes and resuspended and incubated in ACK Lysing Buffer (ThemorFisher Scientific, cat. no. A1049201) for red blood cell (RBC) lysis for 30 seconds at room temperature. RBC lysis was quenched by adding 12 ml cold HBSS++, and brain single cell suspensions were pelleted at 300 g at 4°C for 10 minutes, resuspended in HBSS++ supplemented with 10% vol/vol FBS and maintained on ice until CMO labeling. To obtain single cell suspensions from lungs affected by metastases, lungs were first mechanically digested using scalpels and lung homogenates were incubated in DMEM/F-12 medium supplemented with 10% vol/vol FBS, 2 mg/ml Collagenase Type IV (Millipore Sigma, cat. no. C5138), 1% (w/v) and 30 µg/ml Deoxyribonuclease I at 37°C for 20 minutes in constant shaking at 200 rpm. After enzymatic digestion, lung homogenates were filtered through a 70 µm cell strainer, pelleted at 300 g centrifugation for 5 minutes at 4°C and resuspended in 1 ml ACK Lysing buffer for RBC lysis for 30 seconds at room temperature. RBC lysis was quenched by adding cold PBS, and single cell suspensions from metastatic lungs were resuspended in PBS supplemented with 10% vol/vol FBS and maintained on ice until CMO labeling. For obtaining single cell suspensions from metastatic bones, tibiae and femora from both lower limbs were harvested, cleaned from surrounding soft tissues, and epiphysis were severed and placed on a sterilized mortar containing 1 ml bone digestion buffer, consisting of DMEM/F12 supplemented with 10% FBS vol/vol and 1 mg/ml Collagenase Type II (ThermoFisher Scientific, cat. no. 17101015). Bone marrow samples were flushed with ice cold DMEM/F12 supplemented with 10% FBS vol/vol using a 5 ml syringe coupled to a 27G needle, yielding the bone marrow fraction (BMF), which was maintained on ice. Bone diaphysis were placed in the same mortar with epiphysis containing bone digestion buffer, and using a pestle, mineral bone tissues were crushed until homogenization. Mineral bone homogenates were incubated for 30 min at 37°C, constantly shaking at 200 rpm, and then filtered through a 70 µm cell strainer to remove debris. Both bone marrow and mineral bone fractions were pelleted by centrifugation at 300 g for 5 min at 4°C, and pellets were resuspended and incubated in 1 ml ACK RBC lysis buffer at room temperature for 30 seconds to remove red blood cells. Bone-derived single-cell suspensions were resuspended in PBS supplemented with 10% FBS vol/vol, mixed, and maintained on ice until CMO labeling.

#### CMO labeling

Single-cell suspensions resulting from brain, lung, liver and bones were counted using a Countess II automated cell counter (ThermoFisher Scientific), and cells from each organ were labeled with one CMO of the 3’ CellPlex Kit Set A (10X Genomics, cat. no. 1000261) according to the manufacturer’s instructions. Briefly, 2 × 10^6^ cells in ice cold PBS supplemented with 10% FBS were transferred to a 1.5 ml microcentrifuge tube, pelleted at 300 g for 5 min at 4C, resuspended in corresponding 100 µl Cell Multiplexing Oligos and incubated for 5 minutes at room temperature. Cells and CMO mixtures were then washed by adding 1.9 ml chilled PBS + 10% FBS vol/vol, pelleted by centrifugation at 300 g for 5 minutes at 4C, resuspended in 0.5 ml PBS + 10% FBS vol/vol and filtered through a 30 µm strainers into 5 ml conical tubes (Corning, cat. no. 352235) prior to FACS sorting. Cells were constantly kept at 4°C to preserve viability.

#### Cell sorting for separation of cancer cells, proximal stroma, and distal stroma fractions

CMO-labeled, single-cell suspensions were labeled with Sytox Blue (1:1,000, ThermoFisher Scientific, cat. no. S34857) to detect dead cells and sorted according to GFP and mCherry signal into PBS supplemented with 10% FBS vol/vol using either a FACS Aria II or an Aria Fusion cell sorter (both from Becton Dickinson). For brain, lung, and bone tissues, the entirety of the cell suspension resulting from organ dissociations were sorted. For liver tissues, recovery of >1×10^6^ cells per cellular fraction (cancer cells, proximal and distal stroma fractions) was ensured. Cell sorting resulted in 12 samples, corresponding to cancer cell, proximal and distal stroma fractions from brain, lung, liver and bone cell suspensions. These 12 samples were centrifuged at 300 g for 5 min at 4°C, resuspended in chilled PBS + 10% FBS vol/vol, and counted using a Countess II automated counter. Cells from the same cellular fractions but from different organs were pooled at equal cell numbers into one sample to subsequently generate a scRNA-seq library.

#### Droplet generation and scRNA-seq library preparation

Cell pools were processed following the Chromium Next GEM Single Cell 3’ user guide instructions (10X Genomics, CG000388 Rev B). In brief, cells were resuspended in chilled PBS + 10% FBS at a concentration of approximately 1,000 cells/µl and loaded onto the 10X Genomics Chromium Controller for generating single-cell droplets. A total of 14 libraries were generated, encompassing cell pools from the above-mentioned cellular fractions and organs. Multiplexed, single cell gene expression libraries were sequenced on the Illumina HiSeq. Alignment of 3′ end counting libraries from scRNA-seq analyses was completed utilizing 10x Genomics CellRanger v.6.1.2. Each library was aligned to an indexed mm10 genome using cellranger count.

#### scRNA-seq data analysis

scRNA-seq analysis was performed in R (version 4.3.3) using Rstudio (RStudio 2023.12.1+402 “Ocean Storm” Release (4da58325ffcff29d157d9264087d4b1ab27f7204, 2024-01-29) and the Seurat v5 package. Analysis employed is summarized in Extended data 2b. CellRanger outputs from 14 scRNA-seq libraries were loaded into R using the Read10X() function and Seurat objects from each sample were generated using the CreateSeuratObject() function, where genes that were not detected in at least three cells (min.cells = 3) and cells containing fewer than 200 different genes (min.features = 200) were excluded. Metadata containing the following parameters was added to each sample: mouse’, equivalent to each experimental iteration; ‘compartment’, reflecting the sorted metastatic compartment population; and ‘site’, corresponding to the organ of origin. The ‘site’ parameter was conferred by the multiplexing CMO employed for each sample. Seurat objects containing single cell gene expression data from each sample were merged using the merge() function and percentages of mitochondrial genes (percent.mt) expressed in each cell were calculated using the PercentageFeatureSet() function. Low-quality cells were filtered out from Seurat objects utilizing the subset() function, where only cells with nFeature_RNA > 200 and <8,000 and percent.mt < 20 were carried forward in the analysis. After removing low-quality cells, Seurat objects were split by site, and each of the resulting 4 objects were subjected to the following workflow. Each object corresponding to a metastatic site was normalized using the NormalizeData() function, using the ‘LogNormalize’ method and a scale factor of 10,000. After normalization, the top 2,000 most variable genes were calculated using the FindVariableFeatures() function, utilizing the vst selection method. Data were then scaled for dimensionality reduction using the ScaleData() function and linear dimensionality reduction was calculated with the RunPCA() function using the variable features. The ElbowPlot() function was used to assess the number of principal components (PC) for running the FindNeighbors() and the RunUMAP() functions. Clustering was achieved using the FindClusters() function. Differential gene expression was performed using the FindAllMarkers() function in Seurat, which by default applies the Wilcoxon rank-sum test to compare groups. Top marker genes for each cluster were then visualized using DoHeatmap(), and cluster (cell type) identities were manually assigned after examining the top expressed genes in each cluster using the FindAllMarkers() function. This function only returns genes that have a P value below the specified threshold (default = 0.01). Each cell type from each site was then subsetted using the Subset() function and re-analyzed, which enabled the identification of aberrant clusters and cell doublets, which were removed also using Subset(). Once all poor quality cells were identified and removed, Seurat objects from each site were re-merged to create the main Seurat object encompassing all cells (73,051 cells), or carried forward for metastatic site independent analyses (brain metastasis: 15,146 cells; lung metastasis: 11,725 cells; liver metastasis: 28,232 cells; bone metastasis: 17,948 cells). Dot plots showing expression of top marker genes were generated using the DotPlot() function. Relative frequencies of each cell type per organ were retrieved using the ggplot package. For myeloid subclustering, data was subsetted by these cell types: ‘Mono/Macro’, ‘Microglia’, ‘Kupffer cells’, ‘MP’ (Myeloid Progenitors), ‘Alv. macro’ (Alveolar macrophages), ‘Prolif. Mono/Macro’, ‘Prolif. Microglia’, ‘Prolif. Kupffer cells’, and ‘Neutrophils’.

### Differential abundance testing

We used MiloR^31^ to perform differential abundance testing among the various experimental conditions. We first split the samples by organ and built the k-nearest neighbors graph. From that, we defined the neighborhood of a cell as the group of cells connected by an edge to that cell in the overall KNN graph. We can then define the experimental design; in our case we looked primarily at the DA between proximal and distal stroma. We then count how many cells are from each sample for each neighborhood in order to keep track of the variation in cell numbers between replicates for the same experimental condition (distal vs proximal). We then use the ‘calcNhoodDistance’ function to store the distances between nearby neighbors in the Milo object. With all of these pieces in place, we performed DA testing using the ‘testNhoods’ function to test for differences between proximal and distal stroma. From this test, Milo generates a fold-change and corrected p-value for each neighborhood indicating significant differential abundance. We then assigned a grouping method by finding the most abundant cell type within cells in each neighborhood. The most significant neighborhood groups were filtered by edgeR’s quasi-likelihood F-test (QLF test) and the cutoff is *P* < 0.05.

### Cell-to-cell communication inference

We used CellChat^34^ to analyze cell-cell communication between putative cell types due to potential ligand-receptor interactions. We analyzed each organ separately and only analyzed ligand-receptor interactions due to secreted protein signaling or cell-cell contact. We applied the “standard” CellChat workflow using the variance-stabilized gene expression counts, i.e., the log-normalized UMI counts. That is, for each organ, we performed the following steps. First, we used subsetData to subset the full gene expression matrix to only analyze signal ligand genes or signal receptor genes, reducing computation time. Second, we determined significantly expressed signal ligand genes and signal receptor genes (with respect to cell type groups) using identifyOverExpressedGenes, and, consequently, significantly expressed ligand-receptor pairs with respect to cell type pairs using identifyOverExpressedInteractions. Then, CellChat calculated ligand-receptor interaction scores using the trimean (i.e., type = “trimean”) to calculate the average ligand and receptor gene expression in potential sender and receiver cell type groups, as implemented in computeCommunProb. To avoid biasing of cell-cell communication results due to rare cell types, we removed ligand-receptor interactions involving cell type groups containing fewer than ten cells using filterCommunication (setting min.cells = 10). To analyze common ligand-receptor interactions between cancer cells and monocyte-derived macrophages across all organs, we used subsetCommunication to extract the list of statistically significant ligand-receptor interactions, as determined by CellChat’s permutation test applied to cell type labels, where an interaction was denoted as significant if the computed p-value satisfied *P* < 0.05.

### Flow cytometry analysis of immune cell populations in multi-organ metastasis

For validation of discrete cell type relative abundance in distal and proximal stroma populations, flow cytometry was employed. Single cell suspensions from murine metastasized brain and lungs were generated as described above for the generation of scRNA-seq libraries. For liver single cell suspensions, hepatocyte viability was disregarded, so instead of the 2-step perfusion described above, livers were excised and mechanically digested using surgical scalpels, followed by enzymatic digestion by incubation at 37°C in DMEM-F12 medium supplemented with 10% FBS, 1 mg/ml collagenase IV, and 10 μg/ml DNAse I. Digested livers were then filtered through 70 μm cell strainers, and RBC lysis was performed. For bone single-cell suspensions, only bone marrow flushes were obtained as described above and utilized. After single-cell suspensions were generated, unspecific epitope binding was blocked using purified anti-mouse CD16/32 antibodies (BioLegend, cat. No. 101302, clone 93) for 15 minutes at 4°C, followed by incubation with fluorescently-conjugated antibodies for 30 minutes at 4°C. These primary antibodies were used: anti-mouse CD74-AF647 (1:100, Biolegend, cat. No. 151004); anti-mouse Ly6C-PE-Cy7 (1:100, Biolegend, cat. No. 128017); anti-mouse F4/80-BV605 (1:100, Biolegend, cat. No. 123133); anti-mouse CD45-APC-Cy7 (1:100, Biolegend, cat. No. 103116); anti-mouse Ly6G-BV510 (1:100, Biolegend, cat. No. 127633); anti-mouse CD11b-BV650 (1:100, Biolegend, cat. No. 101259). After primary antibody incubation, cells were washed and resuspended in FACS buffer (PBS + 10% FBS). The SYTOX™ Blue Dead Cell Stain (ThermoFisher Scientific, cat. No. S34857) was added prior to data acquisition as a dead cell dye (1:1000). Cells were then analyzed using a LSRFortessa™ X-20 Cell Analyzer (Becton Dickinson) run by a FACS Diva (Becton Dickinson) software. Data was then analyzed using FlowJo. Fluorescent minus one (FMO) controls were used to set up gates.

### Immunofluorescence

Tissues were harvested from mice after systemic perfusion with 5 mM EDTA (ThermoFisher Scientific) in PBS via post-mortem intracardiac injection. Tissues were then washed in PBS and fixed for 8-12h in 10% formalin at 4°C. After fixation, brain, lung, and liver tissues were placed in 30% sucrose for 4-6h at 4°C. Bone tissues were placed in 20% EDTA at 4°C for 24-48h, until mineral tissues softened. Tissues were then rinsed in PBS, embedded in Tissue Plus OCT compound (Fisher) using plastic cryomolds and frozen at -80°C. OCT-embedded tissues were then sectioned using a Leica CM1950 cryostat. 5-8 μm sections were air dried at room temperature for 1 hour, washed 3 times with PBS for 3 minutes per wash, and blocked with BlockAid Blocking Solution (Thermo Fisher) for 1h at room temperature. Tissue sections were then stained with primary antibodies at 4°C overnight, followed by 3 washes of 3 minutes each in PBS + 0.05% Tween-20, and incubation using secondary antibodies for 1 hour at room temperature. The following primary antibodies were used: Chicken anti-GFP (1:500, Abcam, cat. No. ab13970); Rat anti-mCherry (1:100, Invitrogen, cat. No. M11217); Rat anti-F4/80 (1:100, Invitrogen, cat. No. 14-4801-82); Rabbit anti-Iba1 (1:500, Wako, cat. No. 019-19741); Rat anti-CD74 (1:100, Biolegend, cat. No. 151002); Rabbit anti-MIF (1:100, Abcam, cat. No. ab187064). The following secondary antibodies were used: Goat anti-chicken-AF488 (1:200, Abcam, cat. No. ab150169); Goat anti-Rabbit-AF568 (1:200, Invitrogen, cat. No. A-11011); Goat anti-rat-AF647 (1:200, Invitrogen, cat. No. A21247); Goat anti-rat-AF568 (1:200, Invitrogen, cat. No. A-11077). Micrographs were acquired using a Keyence BZ-X710 microscope using 10x, 20x and 40x objectives. Super-resolution images were acquired as Z-stacks with a Zeiss LSM 900 Airyscan 2 microscope and Zen image acquisition software. All images were collected using a 63×1.4 NA Plan-Apo oil objective. Airyscan processing of all channels and z-stack images was performed in Zen software.

### Immunohistochemistry of human breast cancer metastases

101 archived, paraffin-embedded, human breast metastasis tissue samples were collected following approval by the Institutional Review Board under the UCI 17-05 protocol. 5 um tissue sections were used for *in situ* analysis of MIF and CD74 expression using the ImmPress Duet System (Vector Labs) according to manufacturer’s instructions. Sections were baked at 65°C for 1 h, and then re-hydrated by sequential treatment with histoclear, 100%, 95%, and 75% ethanol, and water. Antigen retrieval was then performed by treatment with a citrate buffer at low pH (Sigma-Aldrich) for 20 minutes at 95°C. Tissues were rinsed with PBS and incubated in Bloxall quenching buffer (Vector Labs) for 10 min at room temperature, after which blocking was performed using 2.5% horse serum (Vector Labs) for 20 minutes at room temperature. Specimens were then incubated overnight at 4°C with primary antibodies in 2.5% horse serum. Primary antibodies: mouse anti-CD74 (1:100, Biolegend, cat no: 326802); rabbit anti-MIF (1:200, Abcam, cat no: ab187064). The following day, sections were washed in PBS and incubated at room temperature for 10 min with the ImmPress Duet Reagent (Vector Labs), which contains secondary antibodies. After 2 washes in PBS for 5 min each, tissues were treated with the ImmPACT DAB EqV (Vector Labs) substrate for 2 min to develop CD74 signal. Tissues were washed again twice with PBS for 5 min, and then treated with ImmPACT Vector Red (Vector Labs) for 2 min to develop MIF staining. Sections were rinsed in PBS and treated with tap water for 5 minutes. Samples were treated for 10 min with Meyer’s hematoxylin (Thermo Fisher) to counterstain nuclei. After hematoxylin counterstain, tissues were washed in water for 10 min, and dehydrated by sequential treatment with 70%, 95%, and 100 % ethanol. Specimens were treated with histoclear for 10 min and mounted with Cytoseal XYL (Thermo Fisher) and tissue coverslips. Samples were then assessed under bright light microscopy using a Keyence microscope. MIF and CD74 expression scores were evaluated as follows: 0, no expression; 1, low expression; 2, moderate expression; 3, high expression. MIF scores reflected signal intensity, whereas CD74 scores reflected relative abundance of CD74^+^ cells. Relative frequencies in organ-specific cohorts for MIF and CD74 scores in cancer cells or stroma were quantified.

### In vivo 4-IPP treatment

For MIF-CD74 axis disruption *in vivo*, 4-IPP (MedChemExpress, cat no HY-110063) was first prepared in a vehicle solution consisting of 5% DMSO, 45% polyethylene glycol 300 (PEG300, MedChemExpress, cat no HY-Y0873), and 45% normal saline (0.9% NaCl). FVB mice injected intracardially with VO-PyMT breast cancer cells were administered intraperitoneally with 4-IPP (MedChemExpress, cat no HY-110063) at a dose of 5 mg/kg body weight or a vehicle solution as control on days 3, 4, 5, 7, 9 and 11 post cancer cell injection. 4-IPP treatment did not result in weight changes or distress in animals. On day 12 post cancer cell injection, animals were sacrificed and *ex vivo* BLI imaging in brain, lung, liver and lower limb bones was used to determine multi-organ metastatic burden.

### Mif knockdown in breast cancer cells

*Mif* knockdown in VO-PyMT and Py8119 breast cancer cells was achieved with miR-E lentiviral vectors^62^ expressing shRNA against Mif gene products. Lentiviral vectors were purchased from TransOmic Technologies as bacterial glycerol stocks, and plasmid DNA was obtained using the PureLink HiPure Plasmid MaxiPrep Kit (Thermo Fisher, cat no. K210007) after expanding bacterial cultures. HEK293T cells were used to generate lentiviral particles to then transduce breast cancer cells. To produce lentivirus, we transfected HEK2913T cells with pMD2.g and psPAX2 packaging vectors encoding for viral packaging, and plasmids encoding the shRNAs against Mif using Lipofectamine 2000 (Thermo Fisher, cat no. 11668019). HEK293T supernatants containing lentiviral particles were collected, centrifuged at 300 rpm at 4°C, and filtered through 0.45 μm filters to remove cells and debris. Lentivirus were used to infect target VO-PyMT and Py8119 cells in the presence of 8 μg/ml Polybrene Infection/Transfection Reagent (Millipore Sigma, cat no TR-1003). 48 h after lentiviral infection, cancer cells were treated with 800 μg/ml hygromycin B (Thermo Scientific, cat no. 10687010) for at least 5 days to select for transduced cells. Mif knockdown was validated by qPCR and Western blot by comparing Mif expression in control hairpin-transduced (shControl) and shMif-transduced cells.

### qPCR

RNA was isolated from cells using the Quick RNA MicroPrep Kit (Zymo Research, cat no. R1050) and cDNA was generated using the iScript cDNA Synthesis Kit (Bio-Rad, cat no. 170-8891). Gene expression was analyzed using the PowerUp SYBR Green Master Mix (Applied Biosystems, cat no. A25741) in a Viia 5 Real-Time PCR System (Applied Biosystems). The following primer pairs were used:

#### Mif

Forward: 5’-CCAGAGGGGTTTCTGTCGGA-3’

Reverse: 5’-CACTGCGATGTACTGTGCGG-3’

#### B2m

Forward: 5’-CCTGGTCTTTCTGGTGCTTG -3’

Reverse: 5’-CCGTTCTTCAGCATTTGGAT -3’

### Immunoblotting

Cells were collected with RIPA buffer (Thermo Fisher, 89900) containing protease and phosphatase inhibitor (Thermo Fisher, 78430) using cell scrapers. Protein lysates were incubated with agitation for 45 minutes, followed by centrifugation at 13,000g for 15 minutes. Supernatant was removed and protein concentration was quantified using Pierce BCA Protein Assay Kit (Thermo Fisher, 23225) according to manufacturer’s instructions. 20 µg protein samples were run on a 12% SDS PAGE gel (BioRad, 4568046) and proteins were wet transferred to a PVDF membrane. Membranes were blocked with 5% BSA diluted in TBS containing 0.1% Tween-20 (5% BSA-TBST) for one hour at room temperature. Primary antibodies diluted in 5% BSA-TBST were added to the membranes and incubated at 4°C overnight. Primary antibodies included: rabbit anti-mouse Mif (1:1000, Abcam, ab187064) and mouse anti-mouse B-actin (1:1000, Santa Cruz, sc-47778). The following day, membranes were washed in TBS containing 0.1% Tween-20 (TBST) followed by incubation with secondary antibodies diluted in 5% BSA-TBST for one hour at room temperature. Secondary antibodies included: Goat Anti-Mouse HRP (1:2000, Thermo Fisher, Cat No. 31430) or Goat Anti-Rabbit HRP (1:2000, Thermo Fisher, Cat No. 31460). Membranes were washed again with TBST and imaged. Protein bands were visualized using Thermo Fisher Chemiluminescent Substrate kit (Thermo Fisher, Cat No. 34579) and imaged on a BioRad ChemiDoc system.

### MIF ELISA

We employed the Mouse MIF ELISA Kit (Proteintech, cat. No. KE10027) to quantify MIF protein in cell culture supernatants of VO-PyMT cells, according to manufacturer’s instructions. Briefly, VO-PyMT cells were plated in cell culture dishes and let attach overnight. The following day, cell medium was replaced and cells were incubated for 24 hours. Then cell supernatants were collected and filtered through 0.22 μm filters using syringes to remove dead cells and debris. These supernatants were diluted 1:25 in cell medium. 100 μl of diluted supernatants were added to ELISA wells containing anti-MIF antibodies. To generate a standard curve, MIF protein standards were prepared as indicated by the manufacturer and plated onto separate wells. Diluted samples, standards, and blanks (cell medium) were incubated at 37C for 2 hours. MIF ELISA wells were then washed 4 times with the kit’s buffer, and 100 ul of anti-MIF primary detection antibodies coupled to biotin were added to the wells and incubated for 1 hour at 37C. Primary antibodies were removed by washing 4 times and 100 ul streptavidin-HRP solution was added to the wells and incubated for 40 minutes at 37C, followed by another 4x wash. Then 100 μl of tetramethylbenzidine substrate (TMB) were added onto wells to develop the signal for 15 minutes. 100 μl of stop solution were added to quench color development, and absorbance at 435 nm from each well was immediately acquired using a VICTOR Nivo Multimode Microplate Reader (Perkin Elmer). To quantify MIF concentration, a 4-parametric logistic (4-PL) curve was fitted using the absorbance from MIF standards, and absorbance from supernatant samples were then extrapolated to the 4-PL curve to determine MIF concentration in pg/ml.

### T cell suppression assays

To obtain macrophages for T cell suppression assays, experimental metastases were first generated in 7-10-weeks-old, female, FVB mice by intracardiac injection of VO-sLP cells. At day 12 post-intracardiac injections, metastasized livers were digested into single cell suspensions by mincing using surgical scalpels, followed by incubation for 20 minutes at 37°C in DMEM/F12 medium supplemented with 10% FBS, 2 mg/ml collagenase type IV, and 10 μg/ml DNAse I. Liver homogenates were filtered using 70 um cell strainers and centrifuged at 300 g. Resulting cell pellets were incubated in red blood cell lysis buffer for 1 min at room temperature. Red blood cell lysis was then quenched by resuspension in PBS, and resulting liver single cells were processed using the EasySep Mouse CD11b Positive Selection Kit II (Stemcell Technologies, cat. No. 18970) according to the manufacturer’s instructions. CD11b+ cells were then sorted using a FACS Aria II flow cytometer. For CD74^-^ monocytes and macrophages, live (SytoxBlue-negative), GFP^-^/mCherry^-^/CD45^+^/CD11b^+^/Ly6G^-^/CD74^-^ cells were sorted. For CD74^+^ MAMs, live, GFP^-^/mCherry^+^/CD45^+^/CD11b^+^/Ly6G^-^/F4/80^lo^/CD74^+^ cells were sorted. Sorted macrophages were then co-cultured in 96-well plates with naive T cells, which were isolated from the spleens of 7-10-weeks-old, cancer-naive, female, FVB mice using the EasySep Mouse T Cell Isolation Kit (Stemcell Technologies, cat. No. 19851) according to the manufacturer’s instructions. Prior to plating, T cells were labeled with the CellTrace Violet Cell Proliferation Kit (ThermoFisher Scientific, cat. No. C34557) according to the manufacturer’s instructions. To stimulate T cell proliferation, we employed Mouse T-Activator CD3/CD28 Dynabeads for T-Cell Expansion and Activation (Thermo Scientific, cat. No. 11456D). T cell-macrophage co-cultures were incubated at 37C, 5% CO_2_, and 90-95% humidity for 72 hours. Co-cultures were then incubated with the Zombie NIR™ Fixable Viability Kit at a 1:500 dilution for 15 min on ice, followed by FcBlock for 10 min at 4C, and then stained with the following antibodies: APC anti-mouse CD4 (clone: RM4-5, BioLegend, cat. No. 100516), and PE/Cyanine7 anti-mouse CD8a (clone: 53-6.7, BioLegend, cat. No 100721) for 30 min at 4C, in the dark. Numbers of CD4 and CD8 T cells and CellTrace signal were analyzed by acquiring equal volumes from each well using a High Throughput Sampler (HTS) of a LSRFortessa™ X-20 Cell Analyzer (Becton Dickinson).

### scRNA-seq immune profiling in metastatic livers

To characterize leukocyte populations in response to Mif expression by cancer cells, livers from FVB mice harboring either shControl- or shMif-transduced VO-sLP cells were harvested at day 10 post cancer cell injections and subjected to processing for scRNA-seq analysis. Whole livers were excised from each mouse, minced using razor blades for <1 min, and enzymatically digested with 2 mg/ml collagenase type IV and 10 ug/ml DNAseq I in DMEM-F12 medium supplemented with 10% FBS, for 25 min at 37C. Liver single cell suspensions were then filtered through a 70 um strainer, pelleted by centrifugation at 500 g, and red blood cell lysis was performed using the ACK lysis buffer (Lonza) by gently swirling for 1 min at room temperature. Lysis was quenched by PBS, and then cells were pelleted, resuspended in 10 ml chilled PBS + 2% FBS, and cell viability as well as live cell counts were quantified using a Countess automated cell counter (Thermo Fisher). A total of 8 x10^6^ cells per sample were then processed for magnetic enrichment of CD45^+^ cells. Briefly, the cells were incubated with a biotinylated anti-CD45 antibody (Biolegend, cat. No.103103). After washing, the CD45-biotin-labeled cells were incubated with anti-biotin microbeads (Miltenyi Biotec), and CD45^+^ cells were enriched by using the LS columns and the midiMACS separator system (Miltenyi Biotec). Upon CD45^+^ cell enrichment, cells were sorted as distal stroma (GFP^-^/mCherry^-^), or proximal stroma (GFP^-^/mCherry^+^) using a BD Fusion sorter. Sorted populations were then counted and subjected to CMO labeling (10X Genomics), according to manufacturer’s instructions. A total number of 12 samples were obtained (from n = 3 shControl-injected mice, and n = 3 shMif-injected mice; each separated into distal and proximal leukocytes), CMO-tagged, and subjected to droplet enabled single cell profiling using the Chromium v3.1 Single Cell expression profiling kit with with Feature Barcode technology for Cell Multiplexing (10X Genomics), according to manufacturer’s instructions. Single cell gene expression and multiplexing libraries were sequenced using an Illumina HiSeq. Sequencing data was analyzed as described in the ‘scRNA-seq analysis’ section above. After removal of poor-quality cells, single cell transcriptomes were clustered using Seurat v5, and cell types were determined based on expression of canonical markers. Total cell numbers of each cell type were estimated by first calculating relative frequencies in relation to the total cells in the Seurat object and then extrapolating that to the total number of cells per sample counted after CD45^+^ magnetic enrichment.

### Immunophenotyping of metastatic livers

For flow cytometry analysis of livers harboring cancer cells upon Mif knockdown, whole livers were harvested 10 days after cancer cell inoculations and subjected to mechanical and enzymatic digestion to generate single cell suspensions, as described in the previous section. All liver tissue was harvested from each mouse and processed the same to avoid batch-to-batch variability. After digestion into single cells, cell viability and total number of cells per liver were determined using a Countess (Thermo Fisher). Cell suspensions were then processed for antibody staining. A total of 4 antibody panels were used for these stainings. 2 panels involved cell surface staining only, and 2 panels involved both cell surface as well as intracellular staining. For cell surface-only panels, a total of 1 × 10^6^ cells were collected per sample, transferred into a well of a U-bottom 96-well plate (Corning), pelleted by centrifugation at 500 g for 3 min at 4C, and resuspended in FcBlock (BioLegend) at a 1:50 dilution in 50 μl PBS + 2% FBS. Cells were incubated in FcBlock for 10 min at 4C, and then 50 μl of primary antibodies were added in PBS + 2% FBS and incubated for 30 min at 4C. Following primary antibody binding, cells were washed twice and resuspended in 200 μl PBS + 2% FBS. For panels involving intracellular staining, 1 × 10^6^ cells were added to a well of 96-well U-bottom plates and first stained with the Zombie NIR live/dead fixable dye (BioLegend, 1:500 dilution in PBS) according to manufacturer’s instructions. Cells were then washed once with PBS + 10% FBS and incubated with FcBlock and cell surface antibody mixes as described above. Following cell surface staining, cells were washed once with PBS + 2% FBS, pelleted and resuspended in 200 μl diluted FoxP3 Transcription Factor Fixation/Permeabilization buffer (eBioscience, cat. No. 00-5523-00) for fixation and permeabilization. Cells were incubated in Fixation/Permeabilization buffer for 20 min at room temperature, protected from light, followed by one wash in 1X Permeabilization Buffer (1:10 dilution in distilled water, Invitrogen, cat. No. 00-8333-56). Importantly, fixation and permeabilization completely removes intrinsic GFP and mCherry fluorescence signals from cancer cells and stroma, enabling the use of antibodies coupled to fluorophores overlapping with GFP and mCherry signals. After washing, cells were pelleted and resuspended in 50 μl Permeabilization buffer containing FcBlock (1:50 dilution) to block unspecific binding, and incubated for 10 min at room temperature, protected from light. 50 μl of antibody mixes targeting intracellular epitopes were then added in 50 μl PBS + 2% FBS and incubated for 30 min at room temperature, in the dark. After antibody incubations, cells were washed twice in Permeabilization buffer, resuspended in 200 μl PBS + 2% FBS, and both cells stained with cell surface-only panels and intracellular panels were subjected to flow cytometry analysis. For cell surface-only stains, the SYTOX™ Blue Dead Cell Stain (Invitrogen, S34857) was added to each sample (1:1000 dilution) right before flow cytometric analysis. All panels were analyzed using a BD Fortessa X20 cytometer using the HTS plate reader, where 150 μl per sample were analyzed. FMOs were used as controls to assess true expression signals, and healthy (tumor-naive) liver single cell suspensions were used to determine true cancer cell and proximal stroma signals. The following fluorescently-conjugated antibodies were used in each panel. Panel 1 (cell surface-only): CD11b-BV650 (1:100, BioLegend, cat. No. 101259), CD19-BV785 (1:100, BioLegend, cat. No. 115543), CD3ε-PE-Cy7 (1:100, BioLegend, cat. No. 155621), CD45-AF700 (1:100, BioLegend, cat. No. 147716), CD31-APC (1:100, Invitrogen, cat. No. 17-0311-82), and NK1.1-APC-Cy7 (1:100, BioLegend, cat. No. 108724). Panel 2 (cell surface only): Ly6G-BV510 (1:100, BioLegend, cat. No. 127633), F4/80-BV605 (1:100, BioLegend, cat. No. 123133), CD11b-BV650 (1:100, BioLegend, cat. No. 101259), CD11c-BV785 (1:100, BioLegend, cat. No. 117336), I-A/I-E-PerCP-Cy5.5 (1:500, BioLegend, cat. No. 107625), Ly6C-PE-Cy7 (1:250, BioLegend, cat. No. 128017), CD74-AF647 (1:100, BioLegend, cat. No. 151004), and CD45-APC-Cy7 (1:100, BioLegend, cat. No. 103116). Panel 3 (cell surface and intracellular): cell surface antibodies encompassed Ly6G-BV510 (1:100, BioLegend, cat. No. 127633), F4/80-BV605 (1:100, BioLegend, cat. No. 123133), CD11b-BV650 (1:100, BioLegend, cat. No. 101259), CD11c-BV785 (1:100, BioLegend, cat. No. 117336), I-A/I-E-PerCP-Cy5.5 (1:500, BioLegend, cat. No. 107625), Ly6C-PE-Cy7 (1:250, BioLegend, cat. No. 128017), CD74-AF647 (1:100, BioLegend, cat. No. 151004), and CD45-PE-Cy5 (1:100, BioLegend, cat. No. 103110); and antibodies targeting intracellular epitopes encompassed CD68-BV711 (1:100, BioLegend, cat. No. 137029), and Ki-67-AF488 (1:100, BioLegend, cat. No. 652417). Panel 4 (cell surface and intracellular): cell surface antibodies encompassed Ly108-PacificBlue (1:100, BioLegend, cat. No. 134608), CD366-BV605 (1:100, BioLegend, cat. No. 119721), CD4-BV711 (1:100, BioLegend, cat. No. 100557), CD62L-BV785 (1:100, BioLegend, cat. No. 104440), CD44-PE-Cy7 (1:100, BioLegend, cat. No. 103030), CD8-PE-Dazzle594 (1:100, BioLegend, cat. No. 100762), and CD279-FITC (1:100, BioLegend, cat. No. 135214); and antibodies targeting intracellular epitopes encompassed FoxP3-PE (1:100, Invitrogen, cat. No. 12-5773-82), CD152-APC (1:100, BD BioSciences, cat. No. 564331), and CD3-AF700 (1:100, BioLegend, cat. No. 100216). Flow cytometry data was then analyzed using FlowJo 10.10.0 (Beckton Dickinson).

### Statistical analysis

For survival Kaplan-Meier analyses in brain metastasis patient samples stained with MIF and CD74, statistical differences in survival curves were calculated by Gehan-Breslow-Wilcoxon test. All tests used for each analysis are indicated in the corresponding figure legends.

### Data availability

scRNA-seq raw sequencing data is available for download at ArrayExpress under accession number E-MTAB-16621. Processed Seurat objects are available for download at Zenodo (DOI: 10.5281/zenodo.18326242). Visualization of cluster resolved single-cell gene expression is available at the Lawson Lab website (https://lawsonlab.org/).

## Acknowledgements

Authors are indebted to M. Oakes, S. Chung, Q. Nguyen and V. Ciobanu from the UC Irvine Genomics Research and Technology (GRT) Hub for their technical assistance with single-cell library sequencing; and V. Scarfone and P. Nguyen from the UC Irvine Sue and Bill Gross Stem Cell Research Center Flow Core for their help with FACS. We thank F. Marangoni, D. Ma, T. McMullen, J.G. Gonzalez, N. Wechter, G. Hernandez, J. Williams, R. Davis, N. Pervolarakis, K. Blake, and all members of the Lawson and Kessenbrock labs for insightful and critical discussions of the work presented here; and Z. Werb for kindly providing VO-PyMT cells. This work was supported by the Feodor Lynen Research Fellowship from the Alexander von Humboldt Stiftung (J.I.); the George E. Hewitt Foundation for Medical Research (P.N.); the National Cancer Institute (NCI) under award numbers T32CA009054 (H.S., I.A., A.H.), 5T32CA009054-44 (I.A.), R01CA244519 and R01CA259370 (S.M.), R01CA234496 (K.K.), R01CA237376-01A1 and 1P01CA288662-01A1 (D.A.L.); the National Institutes of Health (NIH) under award numbers 1F30CA306157-01 (I.A.), R01GM152594 (Q.N.), T32NS121727-01 (A.L.), R01GM147741 (K.K.); the National Science Foundation (NSF) under award number DMS 1763272 (Q.N.); M.P. was supported by a California Institute of Regenerative Medicine (CIRM) fellowship; the UCI Achievement Rewards for College Scientists (ARCS) fellowship (P.H.); the UCI Medical Scientist Training Program under award number NIH/NIGMS 1T32GM154637-01 (I.A.); Johnson and Johnson (S.M.); the Emerging Leader Award from the Mark Foundation for Cancer Research (K.K.); an American Cancer Society (ACS) Research Scholar Grant (RSG) under award number 134389-RSG-039-01-DDC (D.A.L.); and Team Michelle (D.A.L. and J.I.). The results shown here are in part based upon data generated by the TCGA Research Network: https://www.cancer.gov/tcga. This work also utilized the infrastructure for high-performance and high-throughput computing, research data storage and analysis, and scientific software tool integration built, operated, and updated by the Research Cyberinfrastructure Center (RCIC) at the University of California, Irvine (UCI). The RCIC provides cluster-based systems, application software, and scalable storage to directly support the UCI research community (https://rcic.uci.edu).

## Author contributions

J.I. conceptualized the work, designed experiments, analyzed the data and wrote the manuscript. D.A.L. and K.K. supervised the research and co-wrote the manuscript. P.N., H.S. and I.A. critically reviewed the manuscript. P.N., H.S., I.A., A.L. and H.A. helped with animal experiments. H.S. conducted immunoblotting. S.K.C. and S.M. assisted with 2-step liver perfusions. A.A.A. and Q.N. guided CellChat analyses. G.P. performed Milo differential abundance testing analyses. S.K., C.G.S., S.F.H., S.M., C.S., Z.E., and P.V.H. helped with *in vitro* experiments. M.P. helped in conceptualization and experimental design. D.F.T., D.M.E., and R.A.E. organized collection and provided human metastasis specimens and clinical annotations. A.S. and A.M. helped with clinical annotations of patient samples. A.L.P.A.S., V.C., K.N. and E.Z. helped with histology. A.G-A. performed confocal imaging. K.T.E. assisted with intracardiac injections. A.H. and I.A. helped with T cell suppression assays. All authors read and consent to the manuscript.

## Figure Legends

**Extended Data Fig. 1:**
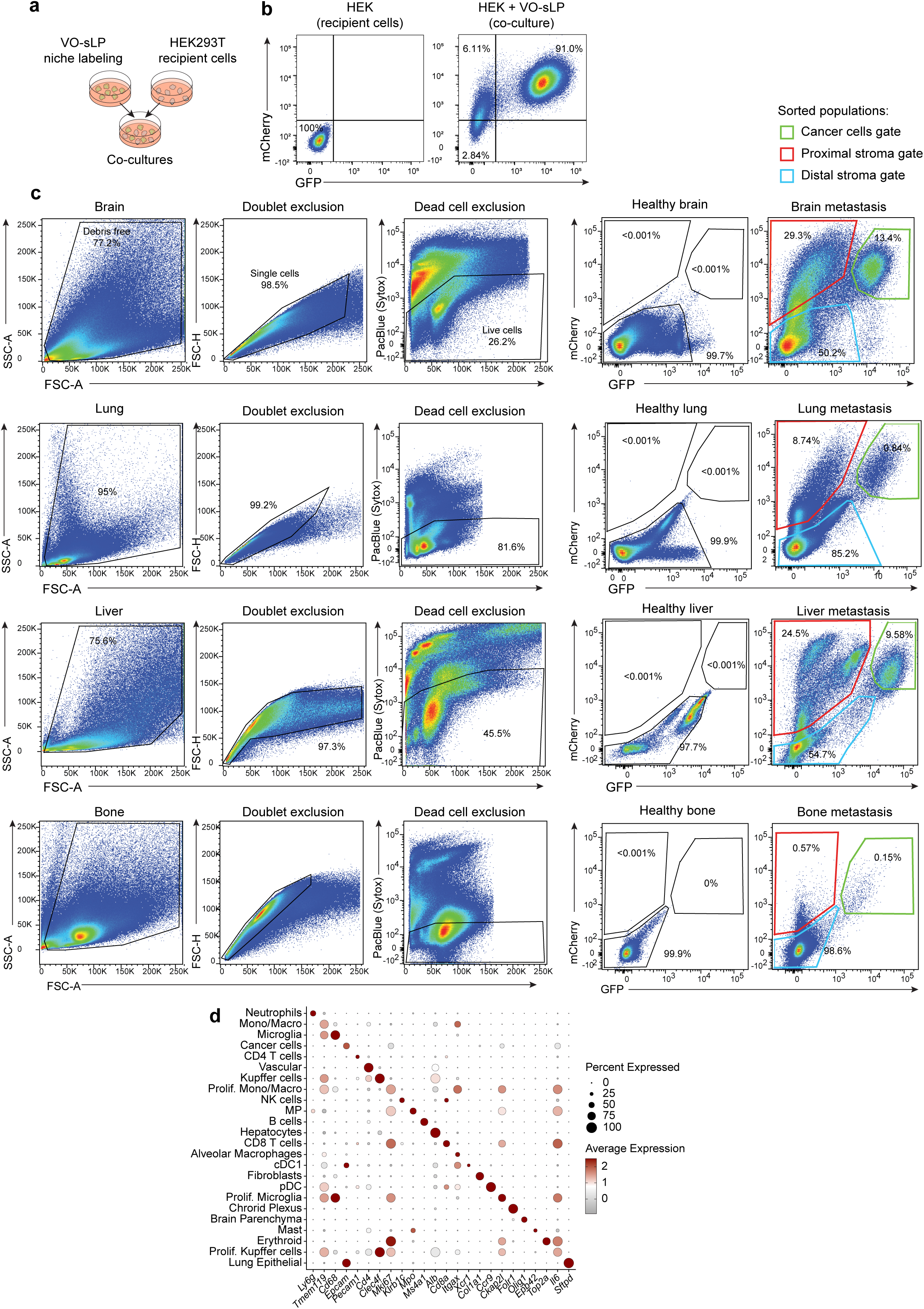
scRNA-seq analysis of niche-labeled multi-organ metastasis. **a.** Schematic depicting *in vitro* validation of niche labeling efficiency in VO-PyMT cells transduced with a GFP/Luciferase construct and the mCherry niche labeling system (VO-sLP)^26,27^. Labeling VO-sLP cells were co-cultured with recipient, non-transduced, HEK293T cells. **b.** Flow cytometry results showing labeling capacity of VO-sLP cells. Plots show GFP and mCherry signals in either monocultures of HEK293T cells (left panel) or co-cultures of VO-sLP cells and HEK293T cells. Cells depicted resulted from excluding cell debris, doublets and non-viable cells (SytoxBlue-positive) by gating using FlowJo. Data is representative of 3 independent experiments. Values inside the plots indicate relative percentages of cells within each gate. **c.** Representative flow cytometry plots showing gating strategy to sort metastatic cellular fractions from brain, lung, liver and bone single-cell suspensions. Data is representative of 3 independent experiments. Numbers within plots indicate relative percentages of each cell population defined by the corresponding gates. **d.** Average expression levels of canonical markers of identified cell types in scRNA-seq data from murine brain, lung, liver, and bone tissues colonized by VO-sLP cells. Dot sizes depict the percentage of cells in each cluster expressing the corresponding gene. Color scale indicates average expression levels. Clusters (cell types) are indicated in the y-axis, while genes are indicated in the x-axis.

**Extended Data Fig. 2:**
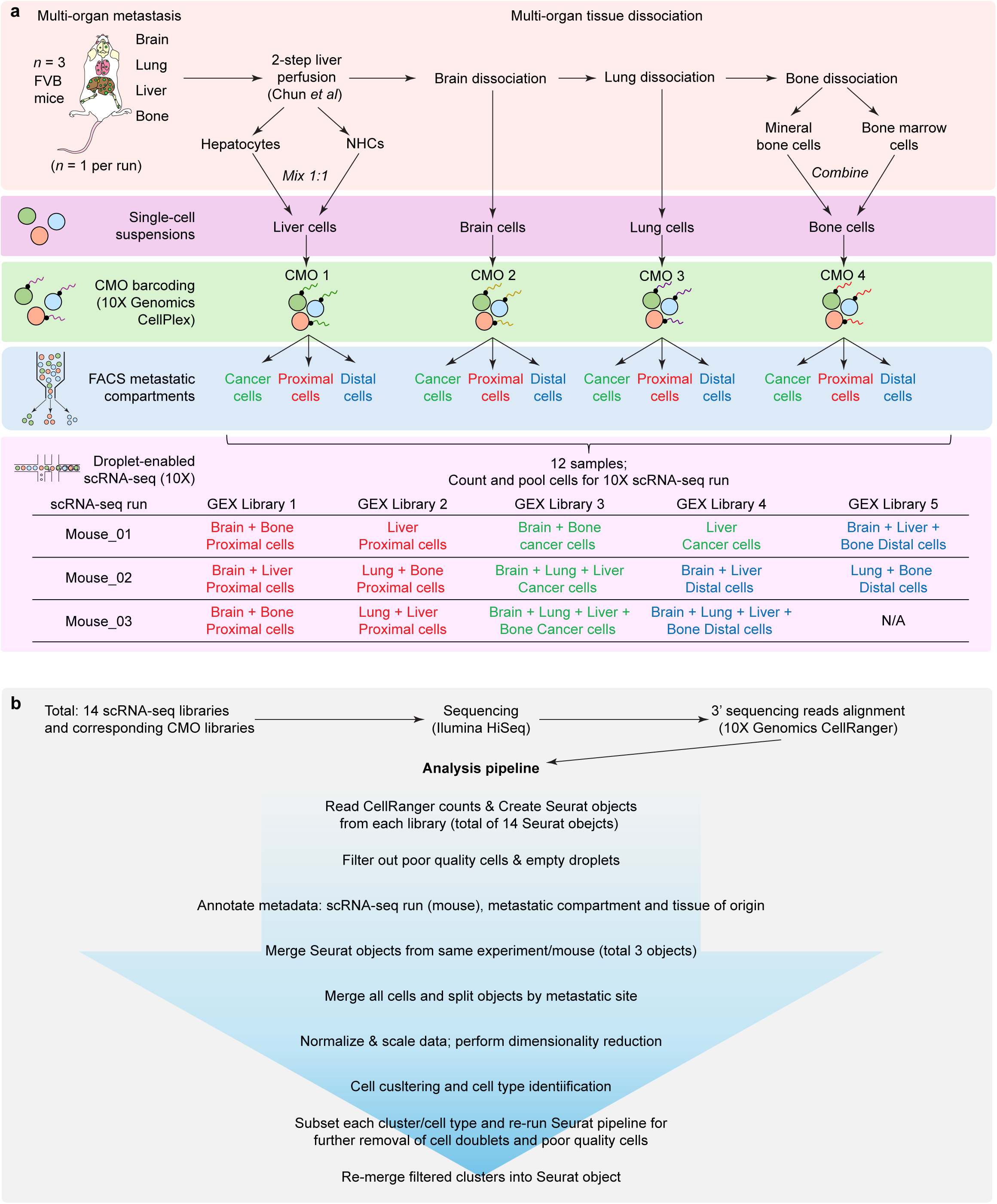
Workflow and analysis pipeline to generate a multi-organ metastasis scRNA-seq dataset. **a.** Overview schematic illustrating niche-labeling scRNAseq approach. FVB mice harboring synchronous VO-sLP-derived metastases in brain, lung, liver and lower limb bones were processed for single-cell preparations, cell sorting of metastatic cellular fractions (cancer cells, proximal and distal stroma), and droplet-enabled scRNA-seq. 2-step liver liver perfusions were performed first at endpoint, followed by brain, lung, and bone tissue dissociation. Resulting single-cell suspensions from each organ were labeled using individual cell multiplexing oligos (CMOs, one CMO per organ) and sorted by flow cytometry according to intracellular GFP and mCherry signal into cancer cells (GFP^+^/mCherry^+^), proximal stroma (GFP^-^/mCherry^+^), and distal stroma (GFP^-^/mCherry^-^) compartments. Upon sorting, CMO-labeled cells were pooled and subjected to droplet-enabled scRNA-seq (10X Genomics). Table shows cellular fractions pooled in each scRNA-seq run and resulting gene expression (GEX) libraries. **b.** Flowchart depicting data analysis workflow. A total of 14 scRNA-seq libraries were obtained and sequenced. Sequencing reads were then aligned using CellRanger (10X Genomics) and analyzed using Seurat. Cell types were manually annotated based on top expressed markers and after filtering out poor-quality cells and empty droplets through several rounds of data cleanup following the pipeline depicted.

**Extended Data Fig. 3:**
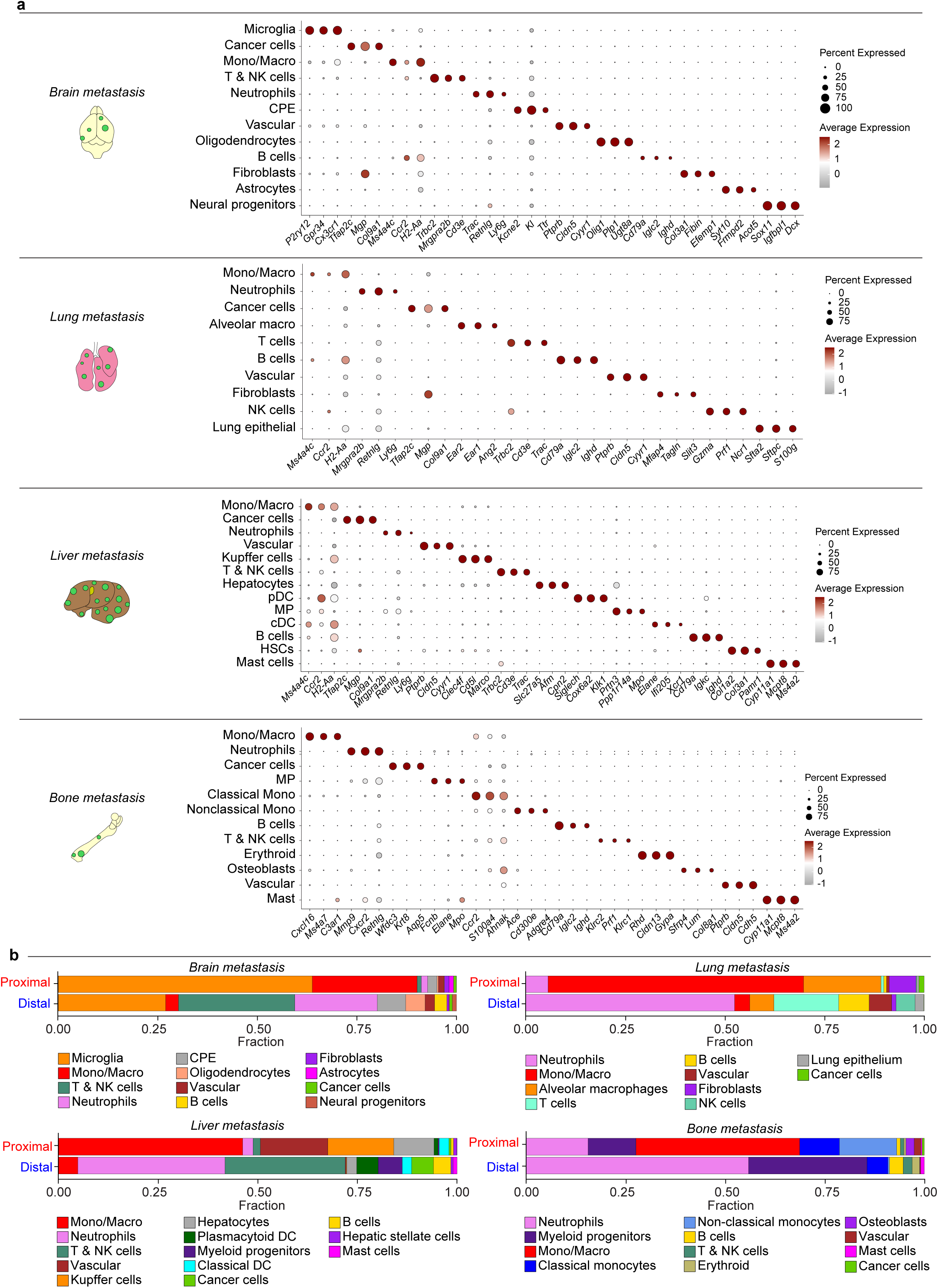
Analysis of metastatic cellular ecosystems in brain, lung, liver and bone. **a.** Dot plots showing expression of 3 of the top markers of each cell type captured in the indicated organs harboring VO-sLP metastases (also see Figure 1). Colors indicate average expression levels and dot sizes depict percentage of cells in each cluster expressing the corresponding gene. Annotated cell types are indicated in y-axes; marker genes are shown in x-axes. **b.** Bar charts showing cell type proportions captured from distal and proximal stroma fractions in metastatic brain, lung, liver, and bone tissues. Relative frequencies of each cell type within distal or proximal populations are depicted.

**Extended Data Fig. 4:**
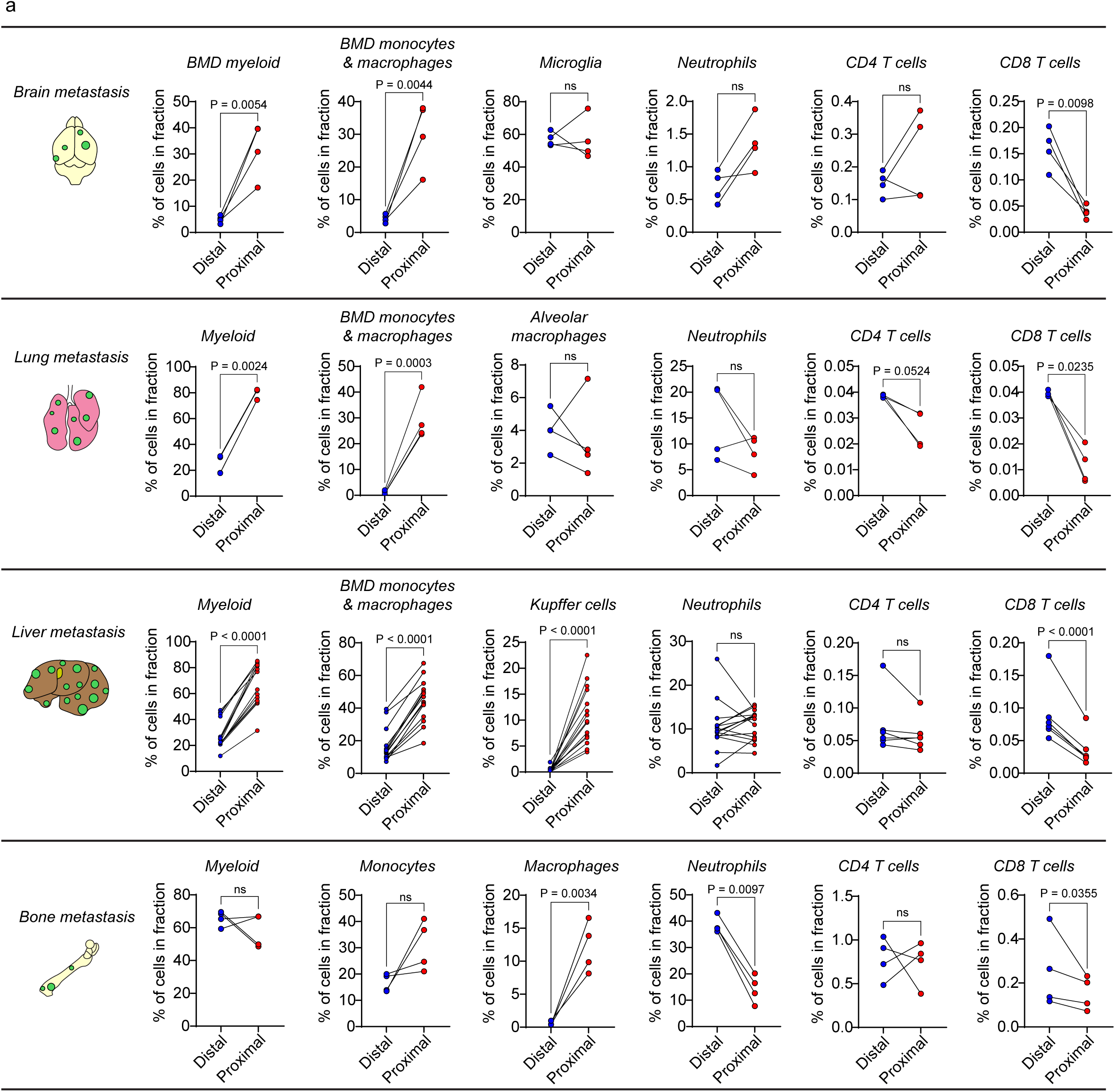
Flow cytometry analysis of myeloid cells and T cells in the proximal and distal stroma of brain, lung, liver, and bone metastasis. Comparison of flow cytometric results between distal and proximal stroma highlighting distinct immune cell populations in the metastatic niche of brain, lung, liver or bone of mice. Cell types analyzed are indicated in the graph titles. Data shows percentages of each population relative to total live cells in either distal or proximal stroma fractions. Dots linked by lines show paired data points from one mouse. *n* = 4; Lung, *n* = 4; Liver, *n* = 14; Bone, *n* = 4. *P* values were calculated using a paired Student’s t test. *ns*, not significant.

**Extended Data Fig. 5:**
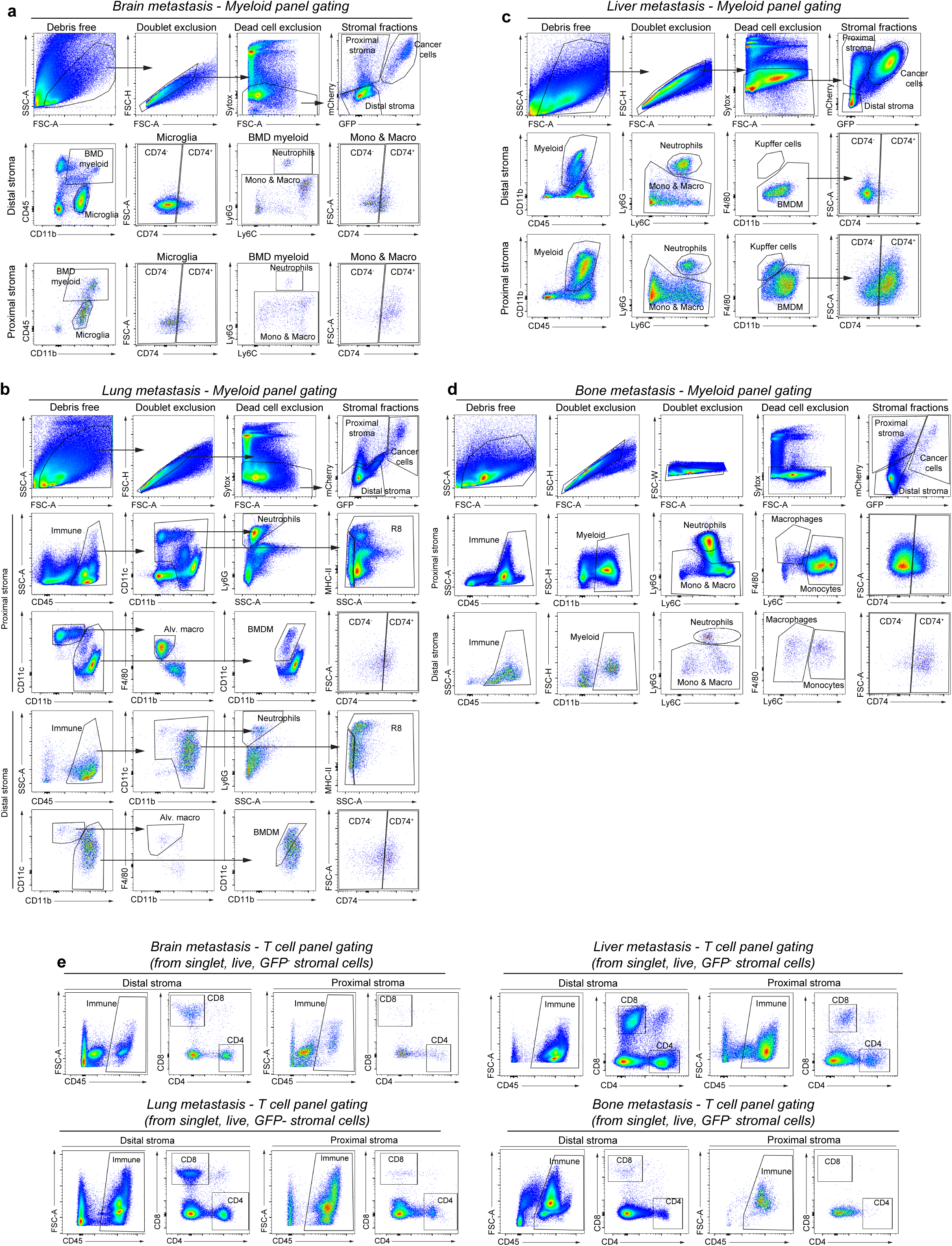
Flow cytometry gating to define immune cell populations in brain, lung, liver and bone metastasis from mice injected with VO-sLP breast cancer cells. **a.** Representative flow cytometry plots showing gating strategy of brain tissues metastasized with VO-sLP cells. Debris free, single, live cells were first separated between proximal (GFP^-^mCherry^+^) or distal (GFP^-^mCherry^-^) fractions. Then, myeloid cells were defined as CD45^hi^CD11b^+^, and microglia were defined as CD45^lo^CD11b^lo^. Myeloid cells were further divided between neutrophils (Ly6G^hi^) and bone marrow-derived macrophages (Ly6G^int/lo^). CD74^-^ and CD74^+^ cells were quantified using the indicated gets. Fluorescent minus one (FMO) stains were used as controls to define the gates for each marker. **b.** Flow cytometry plots showing gating strategy for lungs harboring VO-sLP metastases. For distal and proximal fractions, immune cells were first defined by CD45 signal. CD11c^-^CD11b^-^ cells were gated out, and then neutrophils were defined as Ly6G^hi^. An R8 gate was used to exclude side scatter area-low (SSC-A^lo^) cells. Cells in R8 were then gated for CD11c^hi^, CD11b^lo^, which were used then to define alveolar macrophages (Alv. Macro) as F4/80^+^. CD11b^hi^CD11c^hi^ were defined as bone marrow-derived macrophages (BMDMs). CD74^-^ and CD74^+^ populations within BMDMs were quantified. **c.** Myeloid panel gating in liver samples from mice harboring VO-sLP metastases. In proximal and distal stroma subpopulations, myeloid cells were first defined as CD45^+^CD11b^+^, and then divided in neutrophils (Ly6G^hi^) and monocytes/macrophages (Mono & Macro) (Ly6G^int/lo^). Mono & Macro were further subdivided between Kupffer cells (F4/80^hi^) and bone marrow-derived monocytes and macrophages (BMDMs, F4/80^lo/-^). BMDMs were defined as CD74^-^ or CD74^+^ using the indicated gates. **d.** Representative flow cytometry plots depicting gating for myeloid cells in bone marrow from femora and tibiae colonized by VO-sLP cells. Immune cells were first defined as CD45^+^ cells in proximal and distal stroma fractions. Myeloid cells (CD11b^+^) were divided into neutrophils (Ly6G^hi^) and monocytes/macrophages (Mono & Macro, Ly6G^int/lo^). Cells within the Mono & Macro gate were defined as CD74^-^ or CD74^+^. **e.** Gating in tissue samples harboring VO-sLP metastases. Debris free, single, live cells were divided in proximal (GFP^-^mCherry^+^) and distal (GFP^-^mCherry^-^) stromal subpopulations. For both proximal and distal cells, CD4^+^ and CD8^+^ T cells were quantified within the immune (CD45^+^) subpopulation.

**Extended Data Fig. 6:**
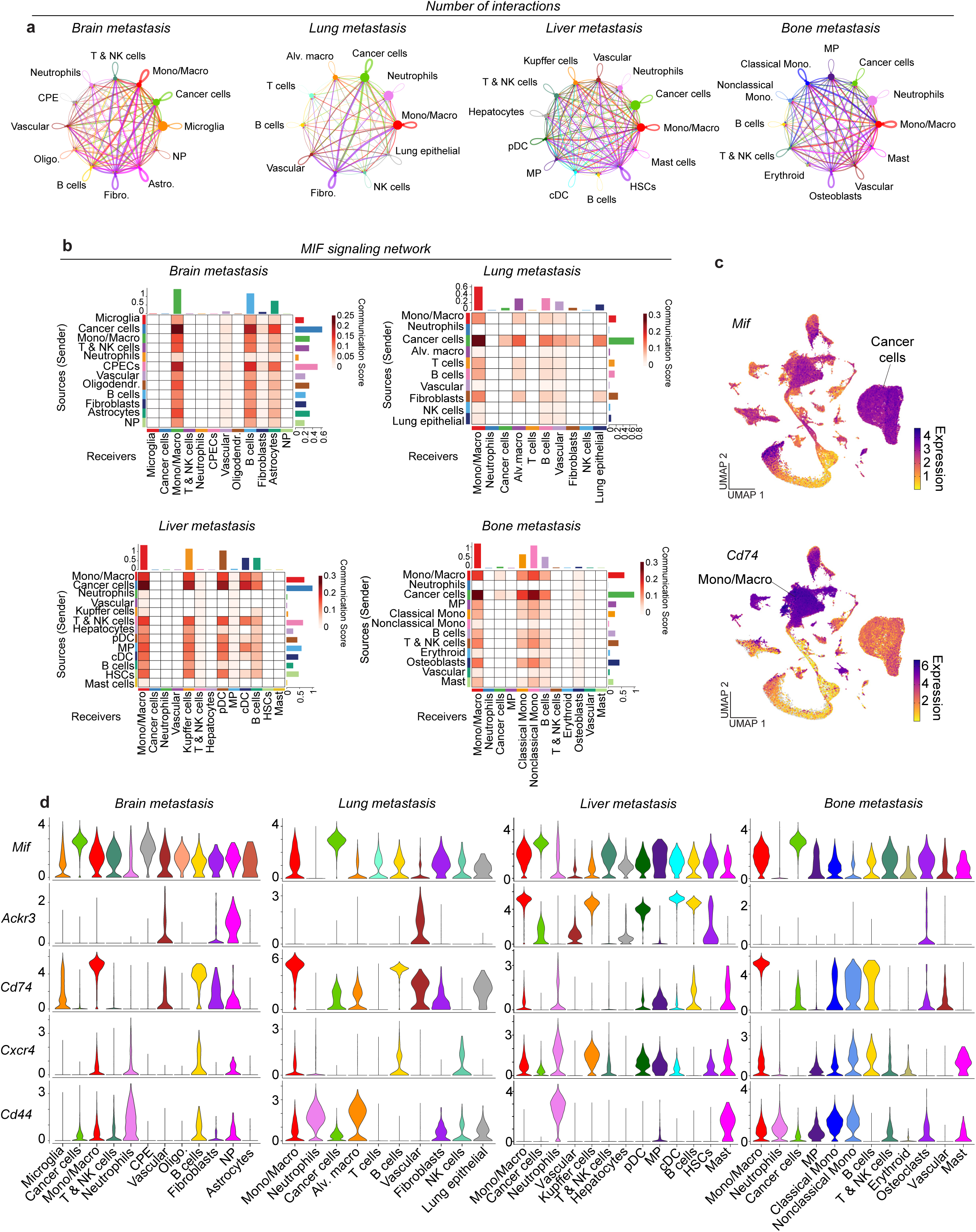
CellChat analysis predicts MIF-CD74 paracrine interaction between cancer cells and MAMs. **a.** Circle plots showing secreted and juxtacrine ligand-receptor interactions between different cell types in brain, lung, liver, and bones colonized by VO-sLP cells, as predicted by CellChat. Arrow thicknesses depict relative numbers of expressed interactions. **b.** Heatmaps showing MIF signaling network communication scores as determined by CellChat analysis between sources of ligands (y-axes) and receptors (x-axes). Barplots at the top of each heatmap show averaged expression of MIF signaling network receptors in the indicated cell types. Barplots in the y-axes on the right side of each panel show averaged expression of MIF ligands in each cell type. **c.** UMAPs depicting *Mif* and *Cd74* RNA expression levels in single cells from FVB mice harboring VO-sLP multi-organ metastasis. Embeddings are as in Figure 1i. Gene expression was analyzed using Seurat. Mono/Macro, monocyte-derived macrophages. **d.** Expression levels of MIF signaling components in metastatic mouse tissues colonized by VO-sLP cells. Data includes expression of the ligand *Mif,* and concomitant components of MIF receptor complexes: *Ackr3, Cd74, Cxcr4,* and *Cd44*. Expression levels of these genes in each cell are depicted by the violins. Y-axes indicate relative expression of the indicated genes; X-axes show cell types. Graph titles indicate the metastatic site analyzed.

**Extended Data Fig. 7:**
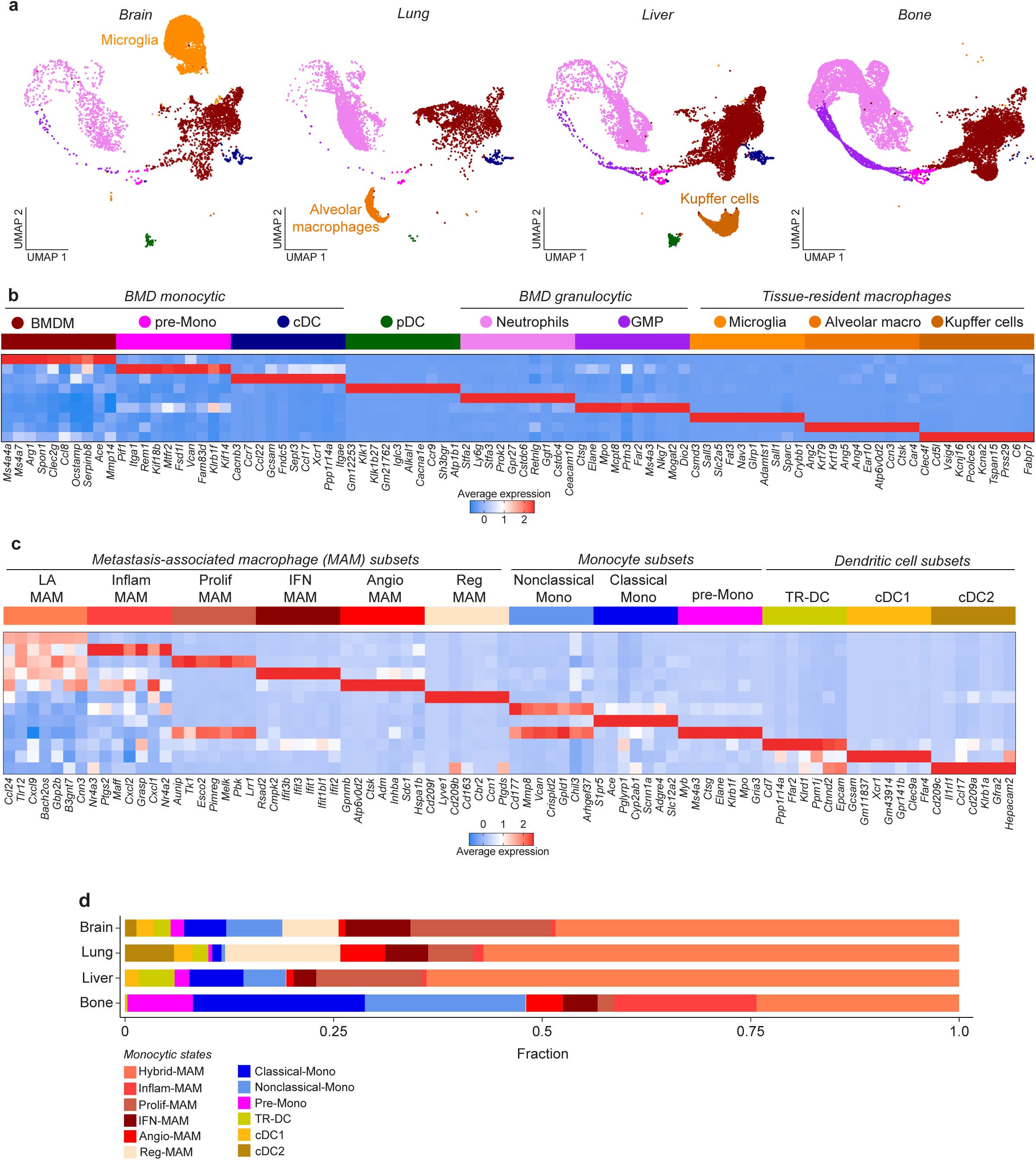
Myeloid populations and monocytic functional states associated with metastasis at multiple sites. **a.** UMAPs showing myeloid cell populations in brain, lung, liver, and bone tissues of FVB mice harboring VO-sLP cells. Tissue-resident macrophages in brain, lung, and liver metastasis are indicated. Cluster colors correspond to labels annotated in (b). **b.** Heatmap showing average expression of top 10 markers for each myeloid cluster as analyzed by Seurat. BMD, bone marrow-derived; Mono/Macro, monocyte-derived macrophages; pre-Mono, monocyte precursors; cDC, classical dendritic cells; pDC, plasmacytoid dendritic cells; GMP, granulocytic myeloid progenitors. **c.** Heatmap depicting top 7 marker expressions for each monocytic cell state, defined by Seurat analysis. Angio-MAM, angiogenic MAMs; cDC1, classical dendritic cells, type 1; cDC2, classical dendritic cells, type 2; Classical-Mono, classical monocytes; IFN-MAM, interferon MAMs; Inflam-MAM, inflammatory MAMs; pre-Mono, monocyte precursors; Prolif-MAM, proliferating MAMs; Reg-MAM, regulatory MAMs; TR-DC, tissue-resident dendritic cells. **d.** Relative frequencies of each monocytic cell state in metastasized brain, lung, liver, and bones, relative to the total number of monocytic cells captured per site.

**Extended Data Fig. 8:**
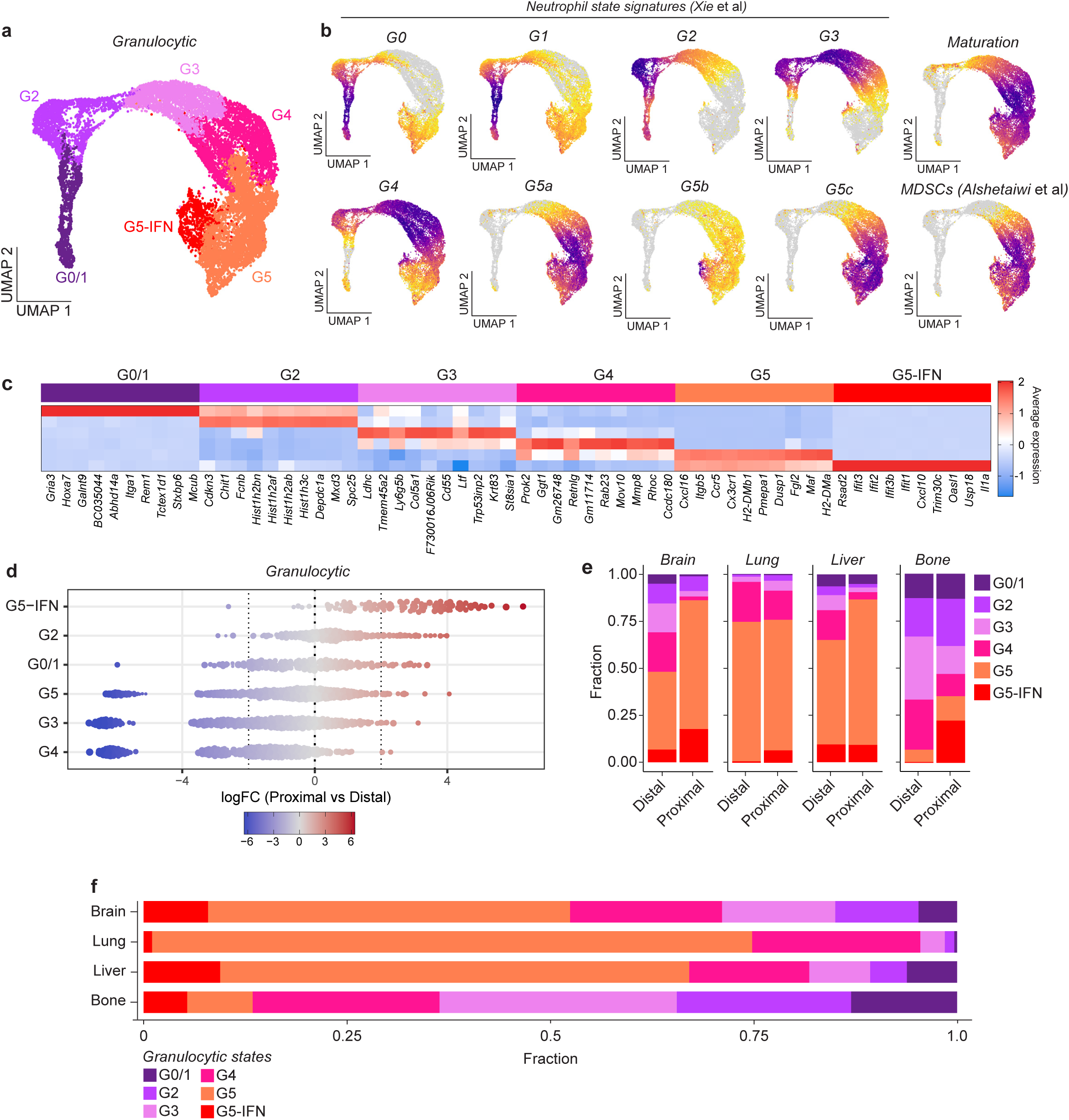
Neutrophil subsets found in multi-organ metastasis. **a.** UMAP showing clustering of granulocytic cells as analyzed by Seurat. G5-IFN, G5 interferon. **b.** Scoring of neutrophil state gene signatures^67^ in granulocytic cells. **c.** Average gene expression of top 10 markers for each granulocytic cell state as identified by Seurat analysis. IFN, interferon. **d.** Differential abundance (log fold change, logFC) of proximal versus distal neighborhoods determined using Milo. Positive logFC values indicate neighborhoods enriched in the proximal stroma (red) while negative values associate neighborhoods with distal stroma (blue). Phenotypic neighborhoods with significant differential abundance (BH-adjusted P ≤ 0.1) are colored according to their corresponding logFC values. **e.** Bar charts showing proportions of different granulocytic cell states in distal and proximal stroma cell fractions at different organs colonized by VO-sLP cells, relative to the total number of granulocytic cells captured in the dataset. **f.** Bar chart showing relative frequencies of granulocytic cell states by organ.

**Extended Data Fig. 9:**
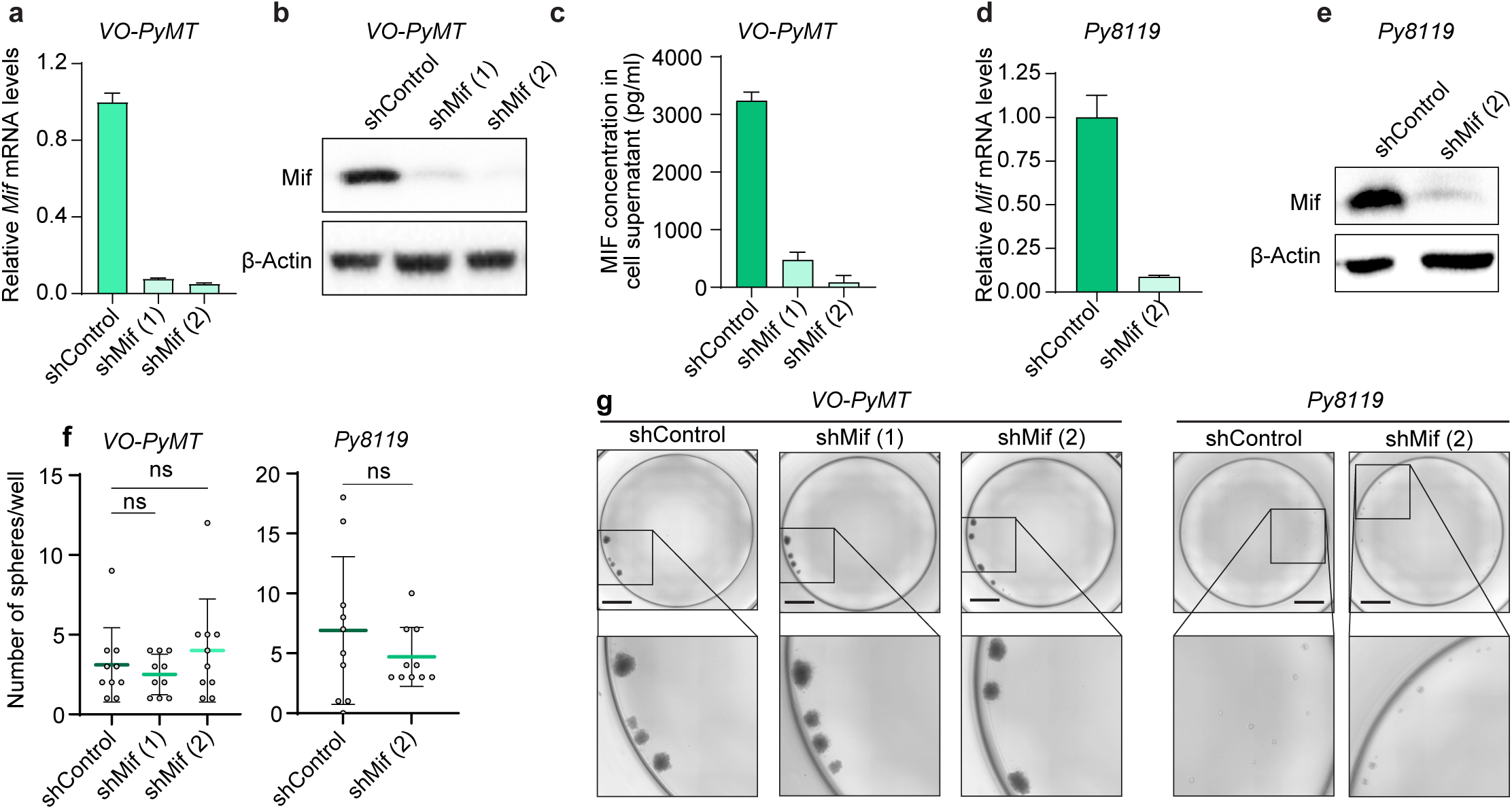
MIF knockdown in VO-PyMT and Py8119 mouse breast cancer cells. **a.** qPCR analysis of *Mif* expression in VO-PyMT cells transduced with control or two independent Mif-targeting short hairpin RNAs (shRNA). Bar heights show means and error bars depict standard deviation of technical triplicates. Data was reproduced in 3 independent experiments. **b.** Immunoblot of Mif protein in shControl- or shMif-transduced VO-PyMT cells. B-Actin is shown as loading control. Data is representative of two independent experiments. **c.** MIF protein concentration in cell culture supernatants of shControl and shMif-transduced VO-PyMT cells as determined by enzyme linked immunosorbent assay (ELISA). Bar heights depict means and error bars show standard deviation of two technical replicates. **d.** Relative Mif expression levels in Py8119 cells harboring an shControl or shMif hairpin, as determined by qPCR. Error bars show standard deviations. Data is representative of 3 separate experiments. **e.** Western blot analysis of MIF protein in shControl or shMif Py8119 cells. B-Actin is shown as loading control. Data is representative of two independent experiments. **f.** Number of spheres per well quantified in *in vitro* spheroid cultures of VO-PyMT or Py8119 cells upon Mif knockdown. Sphere count was performed 7 days post seeding. *Ns,* not significant. Two-tailed Student’s *t* tests were used to estimate significance. **g.** Representative images of spheroids quantified in (f). Scale bars: 1 mm.

**Extended Data Fig. 10:**
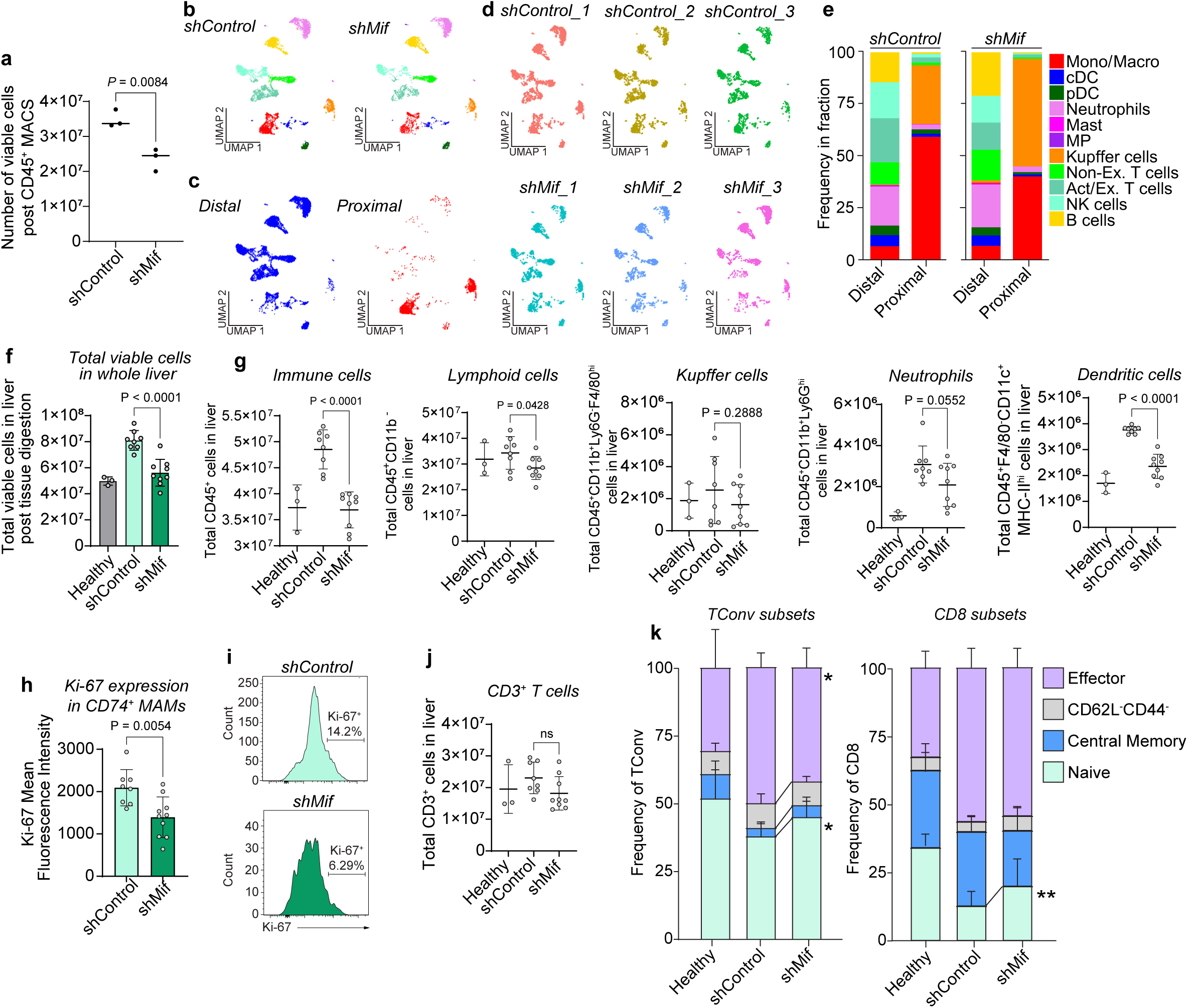
Immunophenotyping analysis of metastatic livers in response to Mif knockdown in cancer cells. **a.** Total number of live cells quantified after processing whole livers for single-cell suspension generation and MACS selection of CD45^+^ cells. Livers were colonized by VO-sLP cells expressing control (shControl) or Mif-targeting (shMif) shRNAs, and harvested at day 7 post cancer cell injections. shControl, *n* = 3; shMif, *n* = 3. Horizontal lines depict median values. Each dot represents data from one mouse. *P* value was determined by a two-tailed Student’s *t* test. **b.** Dimensionality reduction of immune cells from livers harboring either shControl or shMif-expressing VO-sLP metastases, split by group. Embedding of cells is the same as in Figure 6b. **c.** UMAP of metastatic liver immune cells, split by either distal or proximal stroma sorted populations. Embedding is as in Figure 6b. **d.** UMAP split by sample of origin, which corresponds to each mouse used in the scRNA-seq experiment depicted in Figure 6a. UMAP titles indicate identifiers used for each mouse in the experiment. **e.** Relative frequencies of each immune cluster in either shControl or shMif groups. Within each group, frequencies in distal and proximal stroma fractions are compared. N = 3 mice per group. **f.** Total numbers of viable cells quantified after whole-liver dissociations from healthy mice (Healthy), or mice bearing metastatic lesions from VO-sLP cells expressing control (shControl) or Mif-targeting (shMif) shRNAs. Livers were collected 10 days after cancer cell inoculation via i.c. injections. Healthy group, *n* = 3; shControl group, *n* = 8; shMif group, *n* = 9. Bar heights show mean and error bars depict SD. *P* value was determined by a two-tailed Student’s *t* test. **g.** Estimated total numbers of the indicated cell populations in livers colonized by control (shControl) or Mif-deficient (shMif) VO-sLP metastases, or in healthy (tumor-naive) livers. Immunophenotypes of each cell population are indicated in the y-axes. Values were estimated by quantifying the percentage of the population out of total live cells in flow cytometry data, and extrapolating that percentage to the total number of viable cells quantified for each liver after generating single cell suspensions. Horizontal lines show mean values of each group, and error bars depict SD. Healthy, *n* = 3; shControl, *n* = 8; shMif, *n* = 9. Each dot represents data from one whole liver. *P* values were calculated by two-tailed Student’s *t* tests. **h.** Ki-67 expression in CD74^+^ MAMs from livers bearing control and MIF-knockdown VO-sLP cancer cells. Expression levels were measured by mean fluorescence intensity of the Ki-67 signal as analyzed by flow cytometry. Values in each sample result from subtracting mean fluorescence intensity of the FMO control to the mean fluorescence intensity of the sample. shControl, *n* = 8; shMif, *n* = 10. Error bars show SD. *P* value was calculated by a two-tailed Student’s *t* test. **i.** Representative histograms showing Ki-67 expression in CD74^+^ MAMs from data represented in (h). X-axes show Ki-67 fluorescent signals, while y-axes represent cell counts. Segments within plots depict flow cytometry gates used to determine Ki-67^+^ events. Gates were assigned by lack of signal in FMO samples. Numbers indicate percentages of Ki-67^+^ events of the CD74^+^ MAM parent population. **j.** Total CD3^+^ cells estimated in whole liver tissues from healthy mice (*n* =3), or mice injected with VO-sLP expressing shControl (*n* = 8) or shMif (*n* = 9) hairpins. Horizontal lines show mean values in each group. Error bars show SD. Dots depict data from each mouse. *P* values were calculated by two-tailed Student’s t tests. **k.** Relative frequencies of T cell subsets analyzed by flow cytometry. TConv and CD8 populations were defined as follows: Naive, CD62L^+^CD44^-^; Effector, CD62L^-^CD44^+^; Central memory, CD62L^+^CD44^+^. Remaining events were defined as CD62L^-^CD44^-^. Error bars indicate SD of each subset. Asterisks are indicated to show significant changes in the relative frequencies of the adjacent subset between shControl and shMif groups. * *P* <0.05; ** *P* < 0.01. *P* values were determined by two-tailed Student’s *t* tests.

**Extended Data Fig. 11:**
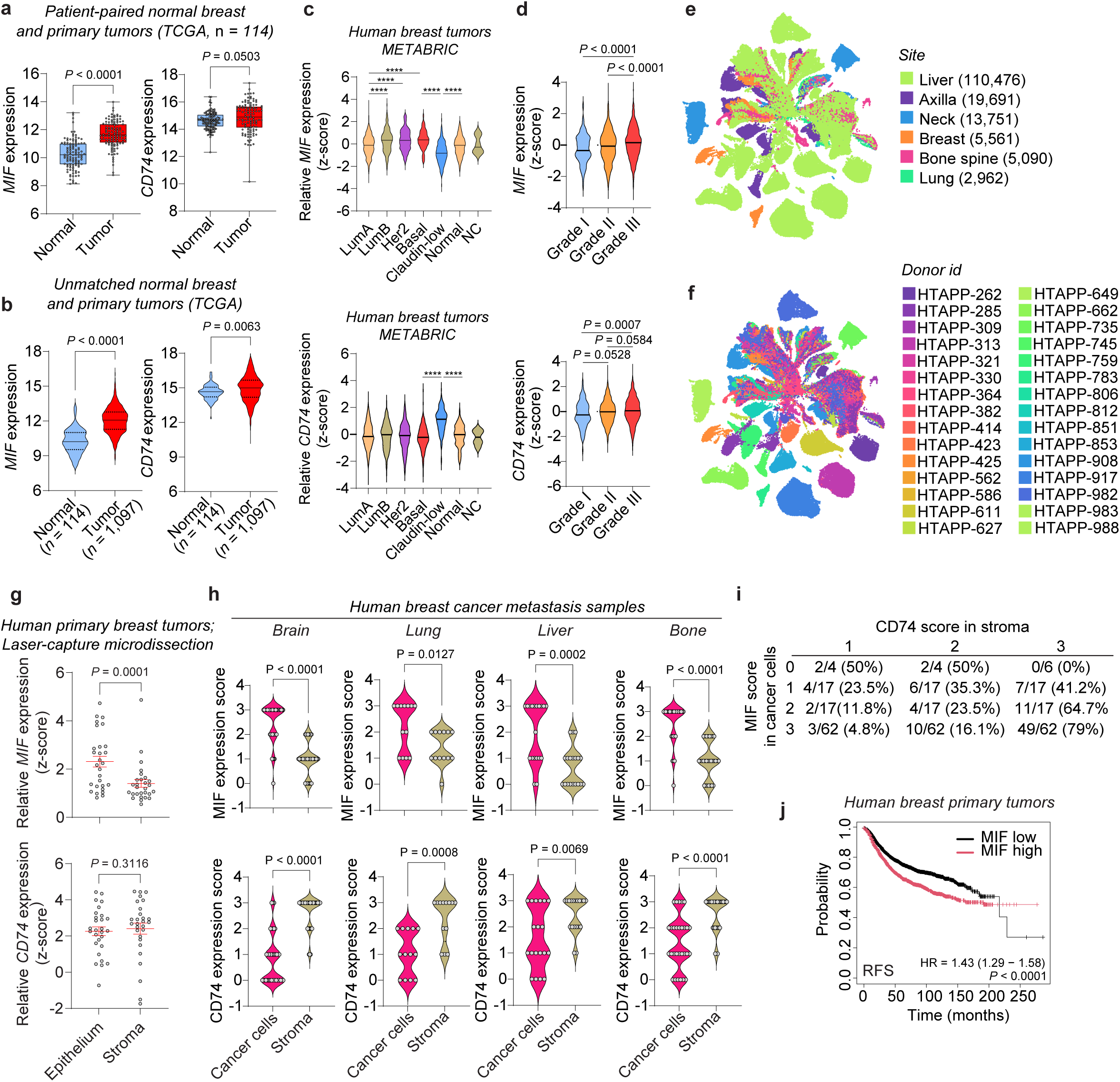
Expression of MIF and CD74 in human breast primary tumors and multi-organ metastasis. **a.** Relative *MIF* or *CD74* mRNA levels in patient-paired normal tissues and primary breast tumors^45^*. n* = 114 patients. Boxes depict inner quartiles. Lines within boxes depict median values. Whiskers show data range, from minimum to maximum values. Each dot represents data from one patient sample. *P* values were calculated with two-tailed paired Student’s t tests. **b.** Comparison of *MIF* and *CD74* relative expression values normal breast tissues (*n* = 114) and breast tumor samples (*n* = 1,097). Violins show the frequency distribution (density) of the data. Median values in each group are represented by straight horizontal lines, while inner quartiles are indicated with dashed horizontal lines within violins. *P* values were determined by unpaired, two-tailed Student’s t tests. **c.** *MIF* and *CD74* expression in a dataset encompassing 1,980 breast primary tumor samples from patients^46^ shown as Z-scores. Graphs show comparison in expression of *MIF* and *CD74* between breast cancer subtypes, as defined by the PAM50 classifier. LumA, Luminal-A; LumB, Luminal-B; Her2, Her2-enriched; Normal, normal-like. NC, non-classified. LumA, *n* = 700; LumB, *n =* 475; Her2, *n* = 224; Basal, *n* = 209; Claudin-low, *n* = 148, Normal, *n* = 148; NC, *n* = 6. **** *P* < 0.0001. *P* values were calculated by ordinary one-way ANOVA usingTukey’s multiple comparisons test. **d.** *MIF* and *CD74* expression in grade 1, grade 2, and grade 3 breast tumors in the same dataset as in (c). Violins show data density distribution and horizontal lines within violins depict median values. Expression values were determined by calculating z-scores for each gene. Grade I, *n* = 169; Grade 2, *n* = 771; Grade 3, *n* = 952. **e.** Tissue-of-origin of cells from a human scRNA-seq dataset^48^ of breast primary tumors and metastases at different anatomical sites. UMAP embedding corresponds to figure 8 a-b. **f.** Dimensionality reduction plot using same embedding as in Figure 8a-b, showing cells colored according to patient of origin (donor id). **g.** *MIF* and *CD74* relative RNA expression levels (Z-scores) in patient-paired epithelium and stroma compartments of breast tumors obtained by laser-capture microdissection (GSE10797)^49^. *n* = 28 patient samples. Horizontal lines show mean values and error bars depict SEM in each group. Each dot represents one data point. *P* values were obtained by two-tailed paired Student’s *t* tests. **h.** MIF and CD74 protein expression scores in cancer cell and stromal areas as determined by semi-quantitative IHC analysis. Data from each metastatic organ was analyzed separately. Each dot represents data from one patient sample. *P* values were determined by unpaired Student’s *t* test. **i.** Contingency table showing correlation in relative frequencies between MIF IHC score in cancer cells and CD74 IHC scores in stroma of brain, lung, liver, and bone human breast cancer metastases. **j.** Relapse-free survival (RFS) probability of patients expressing either high (MIF-high) or low (MIF-low) RNA levels, as calculated by the Kaplan-Meier Plotter^66^. MIF-high, *n* = 1,917; MIF-low, *n* = 3,012. HR, hazard ratio. **k.** Overall survival probabilities in the METABRIC dataset^46^ in either all patients, and in patients expressing either high or low levels of *CD74* in the primary breast tumors, split according to MIF expression. CD74 cohorts were determined by upper tertile cutoff in CD74 expression, where CD74-high represents the upper tertile, and CD74-low encompasses the two lower tertiles. Survival analysis was calculated by splitting patients according to MIF expression, which was cutoff at the median expression value of all samples in the dataset. HR, hazard ratio. *P* values were determined by Gehan-Breslow-Wilcoxon tests.

**Extended Data Fig. 12:**
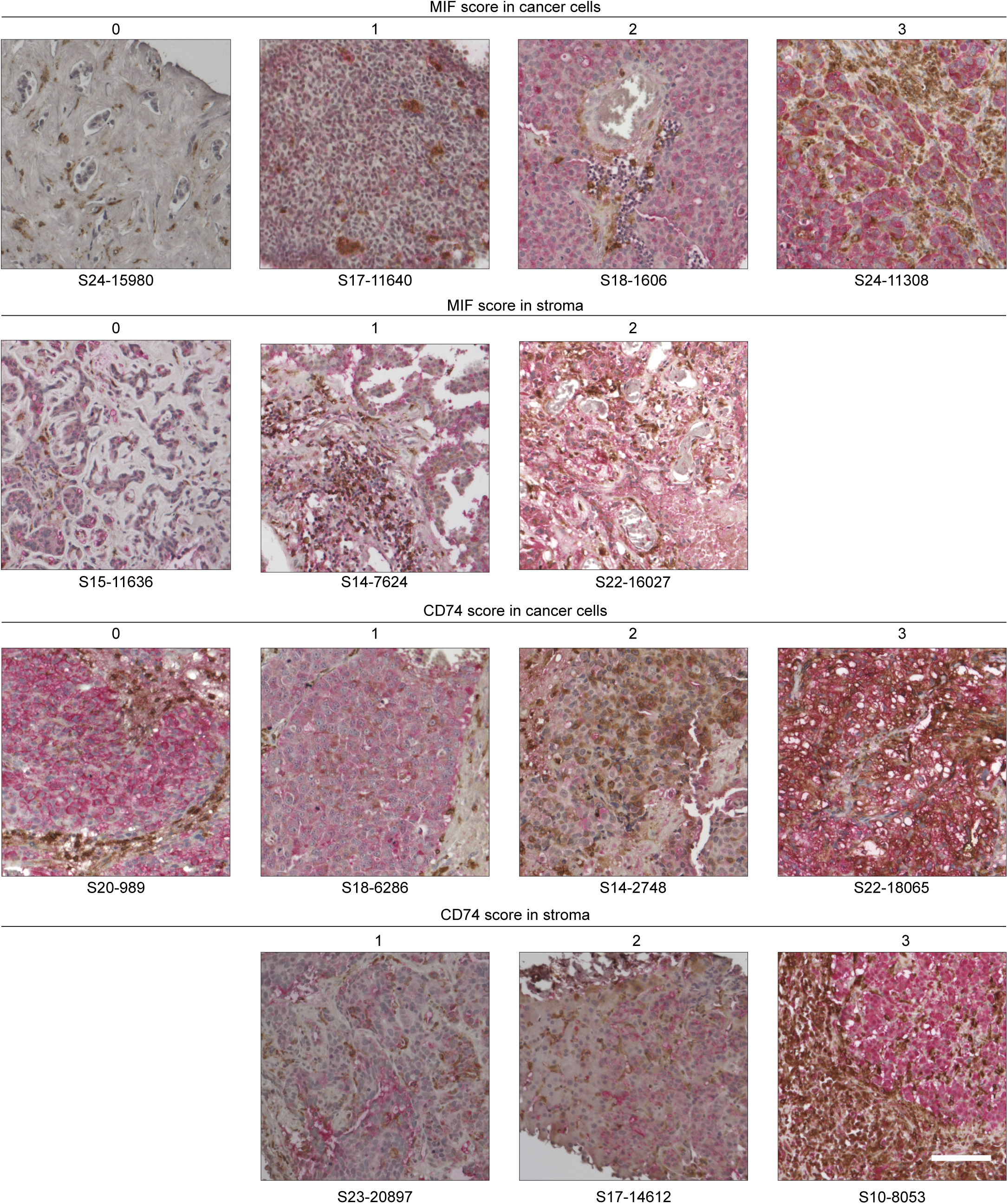
Representative images of MIF-CD74 IHC scoring in cancer cells and stromal areas of human breast cancer metastases. Representative images illustrating semi-quantitative MIF and CD74 scoring approach based on 2-plex immunohistochemistry staining. Scale bars: 100 μm.

## Supplementary Tables

**Supplementary Table 1:** Marker genes associated with each cell type identified in the metastatic niches of the bone, liver, brain, and lung

**Supplementary Table 2:** Cell type marker genes in brain metastasis

**Supplementary Table 3:** Cell type marker genes in lung metastasis

**Supplementary Table 4:** Cell type marker genes in liver metastasis

**Supplementary Table 5:** Cell type marker genes in bone metastasis

**Supplementary Table 6:** CellChat predicted interactions in brain metastasis

**Supplementary Table 7:** CellChat predicted interactions in lung metastasis

**Supplementary Table 8:** CellChat predicted interactions in liver metastasis

**Supplementary Table 9:** CellChat predicted interactions in bone metastasis

**Supplementary Table 10:** Myeloid cell cluster marker genes

**Supplementary Table 11:** Granulocytic cell cluster marker genes

**Supplementary Table 12:** TAM marker genes

**Supplementary Table 13:** Monocytic cell cluster marker genes

**Supplementary Table 14:** Differentially expressed genes in Mono/Macro cells from control (shControl) vs Mif knockdown (shMif) liver tissues

**Supplementary Table 15:** T cell exhaustion signature genes

**Supplementary Table 16:** Clinical metadata for metastatic tumor samples from 100 breast cancer patient cohort

